# Cerebellar acceleration of learning in an evidence-accumulation task

**DOI:** 10.1101/2021.12.23.474034

**Authors:** Marlies Oostland, Mikhail Kislin, Yuhang Chen, Tiffany Chen, Sarah Jo Venditto, Ben Deverett, Samuel S.-H. Wang

**Affiliations:** Neuroscience Institute, Princeton University, Princeton, NJ, USA; Swammerdam Institute for Life Sciences, University of Amsterdam, Amsterdam, the Netherlands; Department of Neurological Surgery, University of California, San Francisco, CA, USA; Department of Anesthesiology, Stanford University Medical Center, Stanford, CA, USA; Present address: Albert Einstein College of Medicine, 1410 Pelham Parkway S, Kennedy 915, Bronx, NY, USA

**Author notes:** Correspondence (S.W.), (M.O.). Equal contribution (Marlies Oostland, Mikhail Kislin).

**Keywords:** Cerebellum, learning, cognition, complex spikes, anterior cingulate cortex, sensory reactivity, behavioral state, task focus, autism

## Abstract

Perturbation to the cerebellum can lead to deficits in motor function, cognition, and behavioral flexibility. Here we report that a cerebellum-specific transgenic mouse autism model with disrupted Purkinje cell function shows unexpectedly accelerated learning on a sensory evidence-accumulation task, as well as enhanced sensory reactivity to touch and auditory cues. Computational latent-state analysis of behavior revealed that accelerated learning was associated with enhanced focus on current over past trials. In on-task states, a subset of Purkinje cells in crus I produced more complex spikes to sensory stimuli. Learning was accelerated by providing cue-locked optogenetic stimulation of Purkinje cells, but unaffected by continuous optogenetic interference with Purkinje cell activity. Complex spikes fired in response to both correct and incorrect choices, but less so when mice were on-task. Both transgenic mice and mice receiving cue-locked optogenetic stimulation showed prolonged sensory responses in Purkinje-cell complex spikes and anterior cingulate cortex. We suggest that cerebellar activity may shape evidence-accumulation learning by enhancing task focus and neocortical processing of current experience.

**Highlights:**

- Faster learning and enhanced sensory salience with cerebellar manipulations in mice
- Accelerated learning arises from prolonged occupancy in an on-task behavioral state
- Cerebellar manipulations can influence neocortex via altered complex spike activity
- Cerebellum findings consistent with a weak global coherence account of autism

**eTOC blurb:** In a cerebellum-based mouse autism model, perturbed function leads to faster learning of a working-memory task, mediated by higher focus on current trials. The effects are emulated by optogenetic perturbation of Purkinje cells, and both perturbations drive enhanced neocortical activity. Results are consistent with a weak coherence model for autism.

## Introduction

The cerebellum’s roles extend beyond movement to include cognition, sensory processing, learning, and memory (Carta et al., 2019; Hatten, 2020; van der Heijden et al., 2021). Recent neuroimaging, clinical, and animal research provides evidence for a cerebellar role in social cognition and adaptive prediction (Frosch et al., 2022; Ito, 2006; Stoodley and Tsai, 2021), and in mice, cerebellar disruption can lead to deficits in attention, behavioral flexibility, and social interaction (Badura et al., 2018). Functional effects can be long-lasting, since early-life cerebellar injury in humans leads to autism spectrum disorder and other nonmotor disabilities (Garfinkle et al., 2020; Küper and Timmann, 2013; Limperopoulos et al., 2007; Tsai et al., 2018; Wang et al., 2014).

Here we present evidence for an enhanced function in cerebellum-specific functional perturbations. To examine alterations in task performance that emerge from abnormal cerebellar circuits, we examined *L7-Tsc1* mutants, a mouse model in which *tuberous sclerosis complex 1* is deleted specifically in cerebellar Purkinje cells (Kloth et al., 2015; Tsai et al., 2012), as well as acute optogenetic perturbations of cerebellar activity. To our surprise, we found accelerated learning on an evidence accumulation task. Further investigations revealed a putative mechanism in which the cerebellum regulates sensory reactivity and salience at brainwide scale to support task persistence and learning.

## Results

*L7-Tsc1* mutant mice have reduced numbers of Purkinje cells (Figure 1A,B), and surviving Purkinje cells show lower firing rates both *ex vivo* (Tsai et al., 2012) and *in vivo* with reduced simple-spike (wild-type mice 55.3±18.8 Hz (mean±SD) in 20 cells from 5 mice, mutant littermates 31.9±14.3 Hz in 34 cells from 4 mice, *t*_(53)_ = 5.06, *p* = 5.5 × 10^-6^) and complex-spike activity (wild-type mice 1.05±0.12 Hz in 20 cells, mutants 0.51±0.19 Hz in 19 cells from 4 mice, *t*_(38)_ = 10.55, *p* = 1.1 × 10^-12^, two-tailed Student’s *t*-test; Figure 1C).

**Figure 1.**
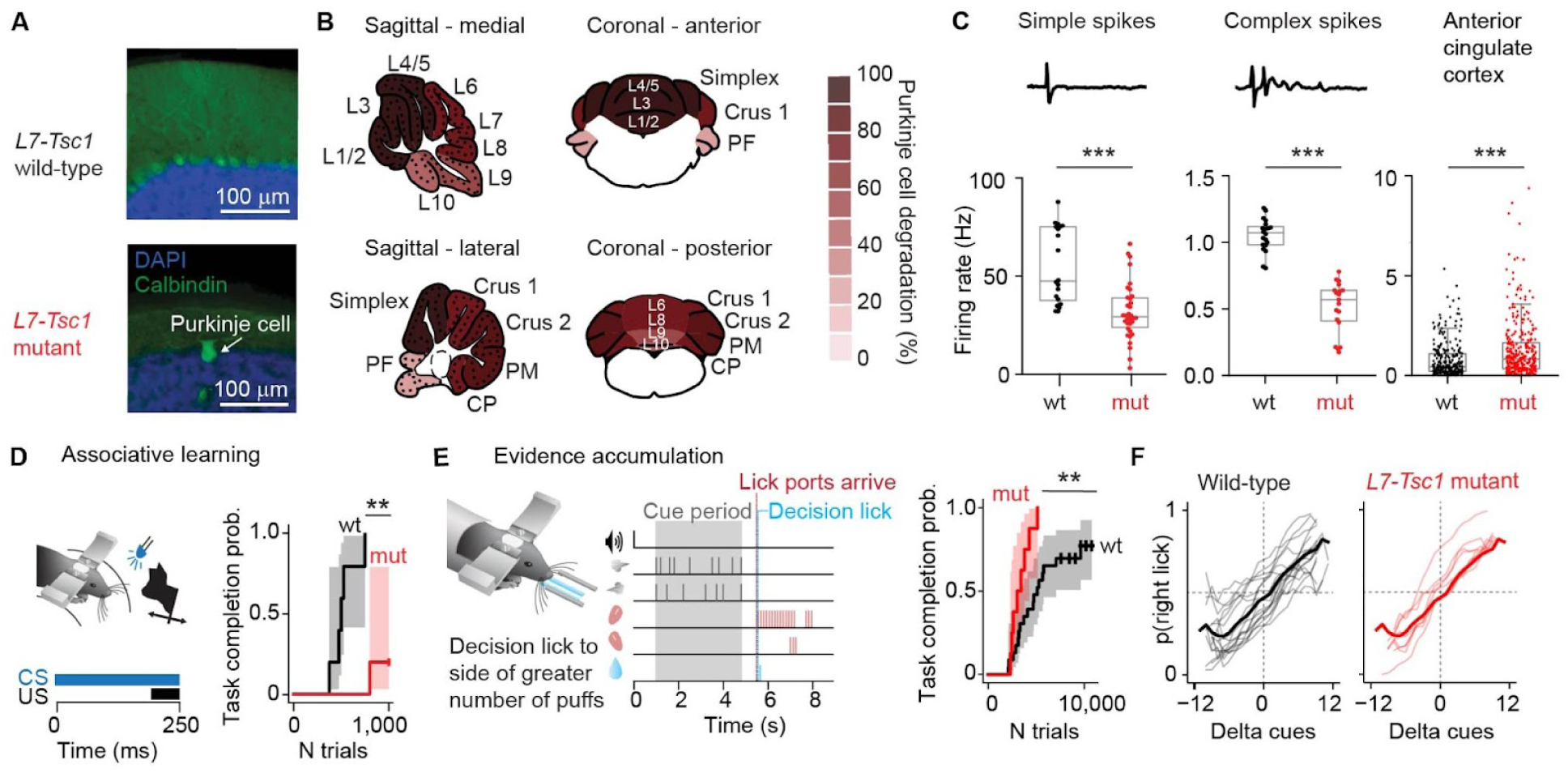
Cerebellar-impaired mice show enhanced learning of an evidence-accumulation decision-making task (A) *L7-Tsc1* mutant mice have reduced numbers of Purkinje cells in the cerebellar cortex. (B) Schematic of sagittal and coronal views of the cerebellum with quantification of Purkinje cell loss averaged over 4 *L7-Tsc1* mutant mice at 5-6 months old for each cerebellar lobule, normalized to 3 wild-type littermates. (C) Reduced spontaneous *in vivo* firing rates of simple spikes and complex spikes, and increased spontaneous *in vivo* firing rates in the anterior cingulate cortex (ACC), in *L7-Tsc1* mutant mice. Example waveforms above the first two plots are 15 ms long. (D) Impaired learning of the delayed tactile startle conditioning task for *L7-Tsc1* mutant mice based on percentage backward CRs. (E) Left: the evidence-accumulation task. Mice receive sensory airpuffs on the left and right whiskers, and receive a reward for correctly licking in the direction of more puffs. Right: Kaplan-Meier estimator of probability of reaching the final level of task training for *L7-Tsc1* mutant mice. Shaded areas in Kaplan-Meier curves represent 95% confidence intervals. (F) Psychometric performance curves in mice who recently reached the final level show no detectable change in bias, slope, or lapse rate. Thick lines indicate averages; thin lines are data from individual animals. Data from all eight *L7-Tsc1* mutant mice are included, and 17 out of 23 control mice (six control animals did not reach the final version of the task).

Despite having deletions of *Tsc1* specifically in the cerebellum, *L7-Tsc1* mutant mice also had higher *in vivo* firing rates in the anterior cingulate cortex of awake behaving mice (Figure 1C; wild-type mean 0.84 Hz in 312 cells from 5 mice, mutant mean 1.34 Hz in 262 cells from 4 mice; *t*_(573)_ = 3.6, *p* = 0.0004), indicative of effects distal to the locus of original perturbation. This matches our past finding that acute chemogenetic suppression of crus I Purkinje cell activity leads to increased activity in anterior cingulate cortex (Verpeut et al., 2023).

*L7-Tsc1* mutant mice show perseveration and deficits in gait and social interactions, as well as deficits in relatively simple motor learning on the accelerating rotarod (Kloth et al., 2015; Tsai et al., 2012). We found that mutant mice were slower to learn delayed tactile startle conditioning behavior (DTSC) (Yamada et al., 2019, Broussard et al., 2022) (mutant mice, median 1000 trials to criterion in 5 mice compared with wild-type littermates 500 trials in 5 mice, *χ*^2^_(1)_ = 9.70, *p* = 0.0018, log-rank test), a cerebellum-dependent form of associative sensorimotor conditioning (Chen et al., 2022). As a means of monitoring progress in DTSC we graphed the time course of learning as a Kaplan-Meier plot (Figure 1D), a method for displaying and analyzing event-occurrence data that allowed us to use tick marks to display mice that failed to complete training by a particular point in time (“censoring” events; Kaplan and Meier, 1958). After 7 sessions, *L7-Tsc1* mutant mice had fewer trials with a conditioned response (CR) than wildtype littermates (wildtype: CR in 62±12 % of trials, mutant mice: CR in 28±16 % of trials, *t*_(9)_ = 3.4, *p* = 0.009, two-sided Student’s t-test).

### *L7-Tsc1* mutant mice have enhanced learning capabilities

We examined a more complex form of cerebellum-dependent learning and information processing by training mice to integrate sensory evidence in working memory using an established evidence-accumulation decision-making paradigm (Deverett et al., 2018; Pinto et al., 2018). Post-learning performance of this task depends on cerebellar crus I (Deverett et al., 2018, 2019), a region that is also necessary for other nonmotor functions (Badura et al., 2018) including visuomotor reinforcement learning (Sendhilnathan et al., 2024) and where *L7-Tsc1* mutants have reductions in Purkinje cells (Figure 1B). During the task, mice receive sensory airpuffs on the left and right whiskers and receive a reward for correctly licking in the direction of more puffs (Figure 1E, see Supplementary Movie S1). Mice progress through increasingly complex levels of task shaping: puffs come to be presented on both left and right, then the difference between the number of puffs on each side becomes smaller, and then an increasing temporal delay separates the end of sensory information from the decision lick (Table S1). As expected, the fraction of correct trials dips when animals transition to the next level of the task, and not all animals reach the final stage (see tick marks on data curves in Figure 1E and 5B, as well as Figure S1A), consistent with the task becoming increasingly difficult with advancing levels.

We were surprised to find that *L7-Tsc1* mutant mice showed *enhanced* learning capabilities, successfully reaching the final level of training almost twice as quickly as wild-type littermates (Figure 1E, mutant median 3211 trials in 8 mice compared with wild-type median 5106 trials in 23 mice; *χ*^2^_(1)_ = 7.11, *p* = 0.007, log-rank test; for individual mouse learning curves see Figure S1A). Acceleration of learning did not vary detectably across training level (number of trials to reach criterion, two-way ANOVA, level × genotype interaction, F_(2)_ = 1.23, *p* = 0.30, comparing early training levels (0, 1, and 2) vs. middle (3 and 4) vs. late training levels (5 and 6)); see also Figure S1B). Learning can depend on sex, age, or stress level (D’Hooge and De Deyn, 2001), but in our case there was no detectable difference in the number of trials by sex, age at start of training, or corticosterone level (Figure S2; F_(3, 18)_ = 0.29, *p* = 0.83, MANOVA).

Once animals had reached an expert stage (level 7), in the first few sessions there was no difference in overall performance between *L7-Tsc1* mutant mice and their wild-type littermates (Figure 1F; no significant difference in percentage correct trials: *p* = 0.53, Welch’s *t*-test, one-tailed; no significant difference in the bias (*t*_(24)_ = 0.81, *p* = 0.43), slope (*t*_(24)_ = 0.10, *p* = 0.92), or lapse rate (*t*_(24)_ = 1.35, *p* = 0.19, two-sided Student’s *t*-tests) of the psychometric curves). Mutant mice also did not differ from wild-types in the number of licks per trial, either in correct trials (*p* = 0.45, two-tailed *t*-test) or incorrect trials (*p* = 0.23). In short, final performance was comparable to animals who have been trained extensively at the same or a similar task (Deverett et al., 2018, 2019; Pinto et al., 2018).

### Latent behavioral-state analysis of task learning reveals bouts of high performance and on-task focus

To explore the alterations in behavior associated with accelerated learning, we performed computational latent-state analysis of the learning process throughout training (Figure 2). Latent-state analysis identifies shifts in behavioral response patterns occurring between groups of trials that reveal variation in internal states over time (Ashwood et al., 2021; Bolkan et al., 2022; Calhoun et al., 2019). We fitted trial-by-trial outcomes using generalized linear modeling of a hidden Markov model (GLM-HMM; Figure 2A), trained on a separate, previously-acquired data set and then fitted to the experimental animals (Figure 2B). The fit was done using previously developed methods of machine learning to define an GLM-HMM (Venditto et al. 2024). In our GLM-HMM, within a training session a mouse is assumed to pass among a user-specified number of states with a fixed trial-to-trial probability. In each of these states, the correct/incorrect performance probability is a function of task parameters (here, current cues, previous trials, and bias). To attain out-of-sample independence for the current behavioral data set, we used a training set of nearly 98,000 trials from 22 wild-type mice from a previous study (Deverett et al. 2018; 2019) to fit a single set of parameters, including state-to-state transition probabilities and a softmax function for converting parameters to performance probability. The likelihood function of the GLM-HMM to capture mouse behavior reached a plateau for 3 or more states, and we chose a 3-state model for further analysis.

**Figure 2.**
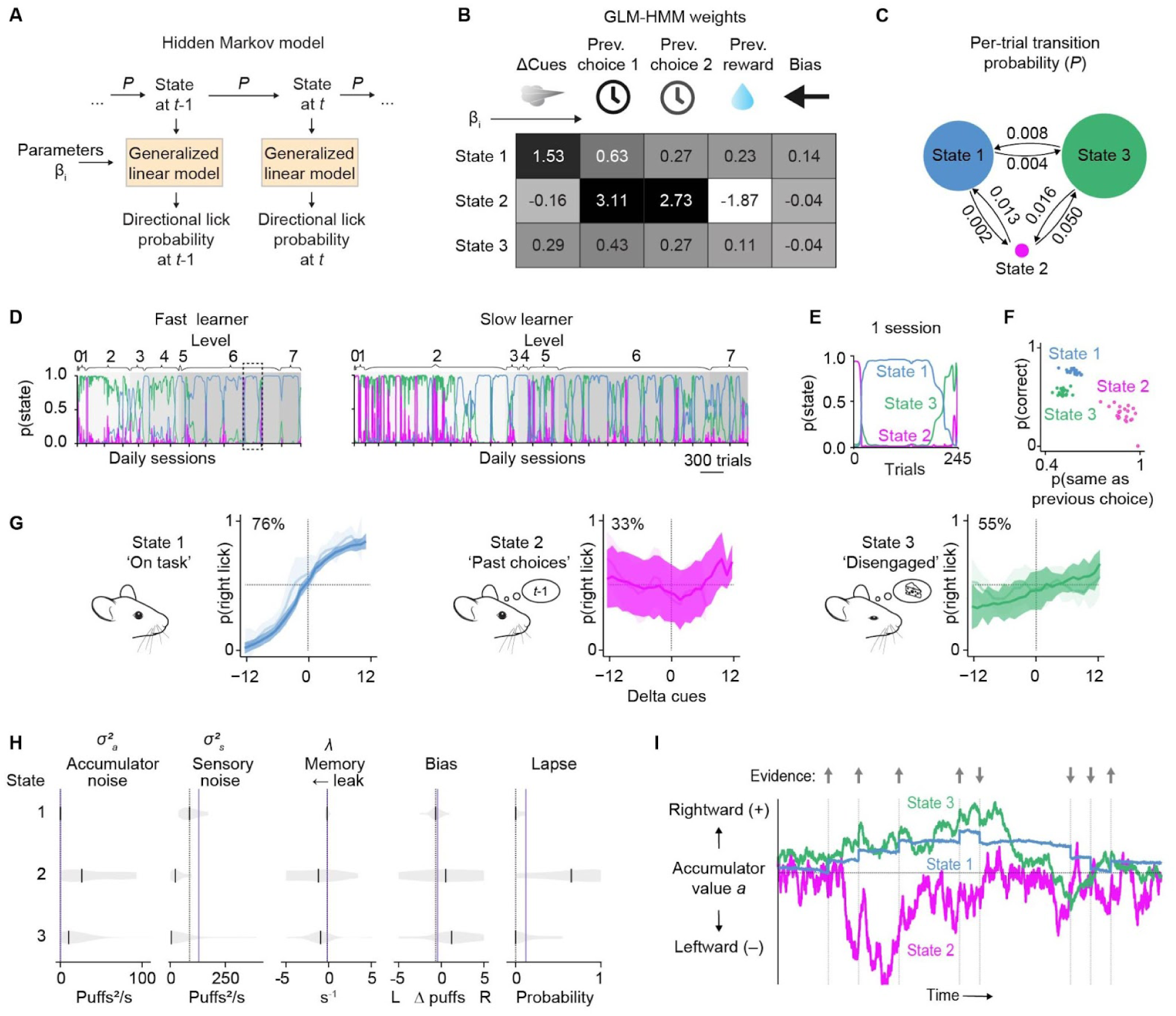
Latent behavioral-state analysis of task learning reveals periods of high performance and on-task focus (A) Schematic of the generalized linear model-hidden Markov model (GLM-HMM). *P* is state transition probability. (B) Inferred GLM-HMM weights from the training data set. (C) Per-trial transition rate between the three states averaged over all mice. The size of the circles indicates state occupancy across all trials of the mice in the training data set. (D) Posterior state probabilities for all trials in all sessions from a fast learner (top) and a slow learner (bottom). The dashed area in the top panel indicates the session in E. (E) Posterior state probabilities for one example session. (F) Probability of a correct choice against the probability that the choice in the current trial was the same as the choice in the previous trial. Each data point represents the average across all trials for one mouse. (G) Psychometric curves averaged across all mice. In the top left is the percentage correct over all trials in that state. Shaded areas represent one s.d. Lighter curves indicate early trials and darker curves indicate late trials. (H) Best-fit drift-diffusion model parameters for the three different behavioral latent states. Fits were computed multiple times for each condition using random subsets of the data to assess the reliability of the best-fit parameters. Black vertical ticks indicate the median best-fit parameter across fit repetitions. Gray shading represents the distribution of fit parameters across repetitions. Vertical purple lines denote best-fit values in one of our earlier datasets (see Deverett et al., 2019). (I) Simulation of the drift-diffusion model. The model’s accumulator value *a* is shown as it evolves over time in a single behavioral trial. Colored lines demonstrate how the trajectory of *a* is qualitatively altered in the three different states. Arrows and associated vertical lines indicate pulses of evidence.

The three states of mouse behavior during the task differed in their dependence on task parameters (Figure 2B). Mice in the on-task state 1 made the most correct decisions, relying heavily on the left-right difference in sensory cues, and less on the animal’s choice in the previous trial. Early in training, wild-type mice tended to spend time in state 2, a past-trial-driven state in which they relied more on past rather than present information, thus reducing their decision accuracy, with responses strongly dependent on the choices made in the previous two trials; and in state 3, an inattentive state in which they made choices but were only weakly sensitive to any evidence-containing features. On a moment-to-moment basis, wild-type mice made transitions from state to state (Figure 2C) on the time scale of dozens or hundreds of trials (Figure 2D). Across sessions as training progressed, animals gradually shifted away from state 2 or 3 occupancy, eventually reaching consistent state 1 occupancy (Figure 2D). This shift in state occupancy occurred across all animals (analysis of variance on the log ratio state 2/state 1, significant effect for training level, F_(2)_ = 53.9, *p* < 0.001; for detailed comparisons, see Figure S3). Within sessions, transitions away from state 1 occurred largely at the end of the session, when animals typically switched from the on-task state 1 to the disengaged state 3 (see example in Figure 2E). Each of these states had a dependency on past trials (Figure 2F) and a distinct psychometric performance curve (Figure 2G) that were consistent with fit parameters.

### In off-task states, accumulated sensory information is noisy and leaky

With different latent states of mouse performance during the task identified, we then sought to understand how the states differed in the processing and retention of sensory evidence. We fitted performance data within each state to a drift-diffusion model we previously used (Deverett et al., 2019) to parametrize the within-trial contributions of sensory noise, accumulator noise, and stability of information storage (Figure 2H, 2I).

We found that state 1’s drift-diffusion parameters were consistent with prior observations in high-performing mice (vertical purple lines replotted from Deverett et al., 2019; Figure 2H). Accumulator noise was low, and the accumulated information was stored with a stable time constant of 5.9±1.1 s (leak rate −0.17±0.03 s^-1^, median±SEM; Supplementary Table 2), comparable to previously observed storage times. In contrast, off-task states 2 and 3 showed high accumulator noise, and accumulated information leaked away rapidly in both state 2 (time constant 0.8±0.6 s, leak rate −1.20±0.88 s^-1^) and state 3 (time constant 1.1±0.4 s, leak rate −0.95±0.34 s^-1^), indicating an emphasis on late-occurring stimuli within each trial. In state 2, the lapse rate was high (0.65±0.12), consistent with the emphasis on irrelevant past trials identified using GLM-HMM analysis. These off-task parameter fits were in a comparable range as previous experiments in which the evidence accumulation and storage process were disrupted by optogenetic interference in crus I (Deverett et al., 2019). Taken together, these findings suggest that whether a mouse is in a state of either excessively attending to past trials (state 2) or disengaging from the task parameters (state 3), it fails to make use of evidence accumulation and storage mechanisms that we have previously shown to require normal cerebellar activity.

### Task-related complex spike responses vary systematically across Purkinje cell dendrites

Generally, complex spikes are evoked by unexpected events to shape action. In the evidence accumulation task, task-related sensory events trigger complex spike activity in crus I during performance during both the cue period and the decision/reward period (Deverett et al., 2018). To test whether complex spike responsiveness varied systematically with behavioral state, we used multiphoton in vivo microscopy (Figure 3A,B,C) to measure calcium transients in 4,224 Purkinje cell dendrites from 3 mice over 13 sessions of training (see Supplementary Movie S2).

**Figure 3.**
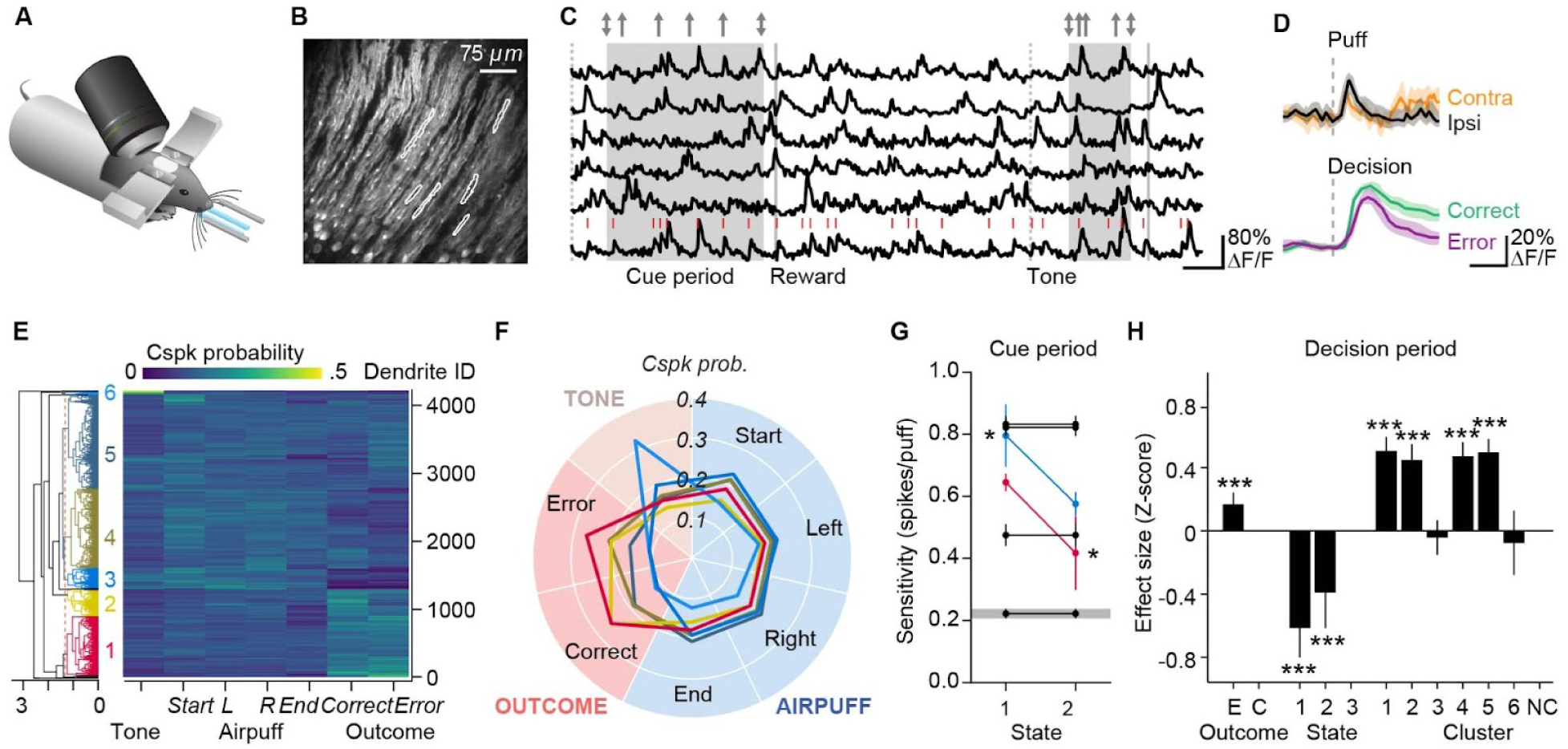
Task-related complex spike responses vary with latent behavioral state (A) Schematic of animal executing the evidence-accumulation task during two-photon imaging. (B) Example two-photon field of view of Purkinje cell dendrites. (C) Signals extracted from dendrites indicated in (B). Dashed lines indicate tone presentation at the start of trials, shaded regions cue periods, arrows at top denote cue timing and side, gray lines mark the timing of decision licks. Red ticks represent dendritic calcium transients extracted from the bottom trace. (D) Top: mean activity of an example dendrite aligned to the moment of individual airpuffs. Bottom: mean response of an example dendritic signal aligned to decision licks. Shading indicates the 95% CI. (E) Left: dendrogram showing hierarchical clustering results for task-related complex spike signals. Right: Heatmap displaying mean response probability of each dendrite at 0-167 ms from the onset of trial events. Cspk = complex spike. (F) Mean task-event tuning for dendrites classified into each cluster in (E). Gray shading indicates the reference level of the linear mixed model. (G) State-dependence of complex spike response per airpuff during the cue period, as measured by the sum of significant main and higher-order effects (post-hoc adjusted means) in a linear mixed model. Asterisks indicate statistically significant three-way interaction effects for state × puffs × cluster. (H) Main effects from a linear mixed model of complex spike activity during the post-decision period. E = error, C = correct outcome, NC = not clustered. Differences are shown using correct outcome, state 3, and unclustered dendrites as reference values.

We assessed dendritic calcium transients in response to task-related events: auditory tone, bilateral puffs at the cue period onset and offset, ipsi-and contralateral airpuffs, and decision lick. Of these events, bilateral puffs at the cue period onset and decision licks showed the most prominent associated calcium transients (Figure 3D). For each imaged dendrite, we calculated the probability of a dendritic calcium transient within a 167 ms window following these events, pooling trials within an imaging session to obtain a within-session probability. Response probabilities were then normalized to a sum of 1 across event types and hierarchically clustered. Clustering revealed six main groups (Figure 3E) comprising 98% of all dendrites which responded most strongly to decision licks (clusters 1 and 2, 21% and 9% of dendrites), ipsilateral puffs (cluster 3, 7% of dendrites), generally to all events (clusters 4 and 5, 28% and 32% of dendrites), and auditory tone (cluster 6, 1% of dendrites) (Figure 3F). The remaining 2% of dendrites were not clustered.

### Latent behavioral state regulates the pattern of evidence representation across dendrites

To examine state-dependent responses to sensory evidence during the cue period, we analyzed 2,740 dendrites that had sufficient trials across behavioral states. We used a linear mixed-effects model (LME, 969,626 observations, AIC = 4.769 × 10^6^) to fit the total number of dendritic calcium transients during the cue period (putative complex spikes, *C_cue_*) to a function of total number of puffs (bilateral and unilateral puffs), categorical variables for state (reference: state 3) and cluster (reference: unclassified dendrites), interaction effects of puffs × state × cluster, with random effects for mouse and dendrite.

The mixed effects model revealed three significant effects. There was a strong main effect of puffs on *C_cue_* (F_(1, 35.64)_ = 206.1, *p<*10^−12^, ANOVA marginal test, Satterthwaite criterion), indicating that the number of puffs strongly influenced neural responses. We also found a highly significant main effect of cluster (F_(6, 247.48)_ = 237.58, *p<*10^−12^), showing that different neural clusters had distinctive baseline activity levels. Finally, a significant puffs × cluster interaction (F_(6, 603.25)_ = 183.42, *p<*10^−12^) demonstrated variation in the responsiveness of different clusters to puffs.

Although state by itself had little or no effect on per-puff sensitivity (state × puffs, F_(2, 10780)_ = 0.66, *p* = 0.518) or baseline activity (F_(2, 9913.6)_ = 2.76, *p* = 0.064), we did observe a highly significant state × cluster interaction (F_(12, 8816.7)_ = 7.12, *p<*10^−12^), as well as a three-way interaction between state, puffs, and cluster (F_(12, 9618.9)_ = 9.34, *p<*10^−12^). Thus the distributions of both baseline and per-puff complex spike activity were regulated across clusters in a behavioral state-dependent manner.

In particular, two interactions with behavioral state affected cue-period responsiveness. Summed statistically-significant main, two-way, and three-way spike-per-puff effects are shown in Figure 3G. The on-task state 1 increased the puff-dependence of cluster 3 from 0.58±0.04 spikes/puff (mean±SD) by an additional 0.23±0.09 spikes/puff (*p* = 0.014) to a total of 0.81±0.10 spikes/puff. Secondly, the past-choice-focused state 2 reduced the puff-dependence of the outcome-dependent cluster 1 from 0.65±0.03 spikes/puff by 0.23±0.12 spikes/puff (*p* = 0.049) to 0.42±0.12 spikes per puff. In short, the current-evidence-focused state 1 showed enhanced representation of current evidence compared with the previous-choice-focused state 2 in specific functional clusters totaling 21+7=28% of dendrites.

### The on-task state suppresses complex-spike responses to trial outcomes

Complex spikes also fired in the 800 ms after the decision lick, a point during head-fixed tasks when a correct/incorrect choice leads to a reward, or not. To obtain fit coefficients in units of effect size, we fitted an LME to Z_decision_, the Z-scored number of dendritic calcium transients during the decision period (1.08±1.05 transients, mean±SD), as a function of behavioral state, dendrite cluster, and error-vs.-correct outcome. We included two-way and three-way interaction effects as well as random effects by mouse and dendrite.

This mixed-effects model revealed that outcome-period firing Z_decision_ varied strongly by state (F_(2, 17.304)_ = 22.33, *p* = 1.6×10⁻⁵, ANOVA marginal test, Satterthwaite criterion), outcome (F_(1, 67.219)_ = 18.273, *p* = 6.2×10⁻⁵), and cluster (F_(6, 26.3)_ = 52.747, *p<*10^−12^). Behavioral state had strong effects on cluster-dependent activity (state × cluster, F_(12, 109.55)_ = 9.179, *p* = 5.2×10⁻¹²; state × cluster × outcome, F_(12, 4843.4)_ = 5.2447, *p* = 7.6×10⁻⁹), revealing that state regulated the outcome-dependent distribution of firing across clusters.

The largest main effects arose from behavioral state. Compared with state 3, the on-task state 1 was associated with a 0.61±0.10 smaller response (*t* = 6.19, *p* = 6.2×10^-10^) and the past-trials state 2 showed a 0.39±0.12 smaller response (*t* = 3.38, *p* = 7.3×10^-4^). These effects were partially counteracted by significant state × cluster effects for clusters 1, 2, and 3 (median increase of 0.27). Every one of the effects was larger than the main effect of decision error, an increase of 0.17±0.04 (*t* = 4.27, *p* = 1.9×10^-5^). Thus the overall trend (Figure 3H) was one in which during the outcome period, Purkinje cells produced most complex spikes in state 3, fewer in state 2, and least in state 1. This trend was strongest in clusters 4 (strong response to cues), 5 (error trials), and 6 (tone).

In error trials, increased complex spike responses were both cluster-specific (cluster × error) and state-specific (state × cluster × error, but not state × error). The tendency of mice to produce more spikes in state 3 was present in 5 out of 6 clusters, the only exception being cluster 3, a puff-sensitive cluster. Overall, state 3 showed a response of 0.43 more than state 1, well over one-third of a standard deviation. In short, the encoding of decision errors via complex spikes was highly state-dependent, and the most spikes occurred when mice were off-task.

### Mutants stay on-task and in the present

To measure how differential occupancy of task states evolves over training, we made use of our division of the behavioral data into three stages of task shaping, early (levels 0-2), middle (levels 3-4), and late (levels 5-6) (Figure 4A). Compared with wild-type mice, *L7-Tsc1* mutant mice had higher state 1 occupancy at later stages of training (Figure 4B). Using compositional analysis of state occupancies, multivariate analysis of variance on the log ratio state 2/state 1 yielded significant results for wild-type vs. *L7-Tsc1* mutant mice (F_(1)_ = 18.8, *p* < 0.001) and for training level (F_(2)_ = 24.5, *p* < 0.001). Post hoc comparisons using the Tukey honestly-significant-difference test indicated a significant result for later stages of training (levels 5-6, *p* = 0.006), but not for early (levels 0-2, *p* = 0.49) or middle stages of training (levels 3-4, *p* = 0.29). Multivariate analysis of variance on the log ratio state 3/state 1 yielded a trend towards significance for wild-type vs. *L7-Tsc1* mutant mice (F_(1)_ = 3.6, *p* = 0.06) and no significant effect for training level (F_(2)_ = 1.9, *p* = 0.15). The shape of the psychometric curves of *L7-Tsc1* mutant mice in each state was similar to their wild-type littermates (Figure S4A,B).

**Figure 4.**
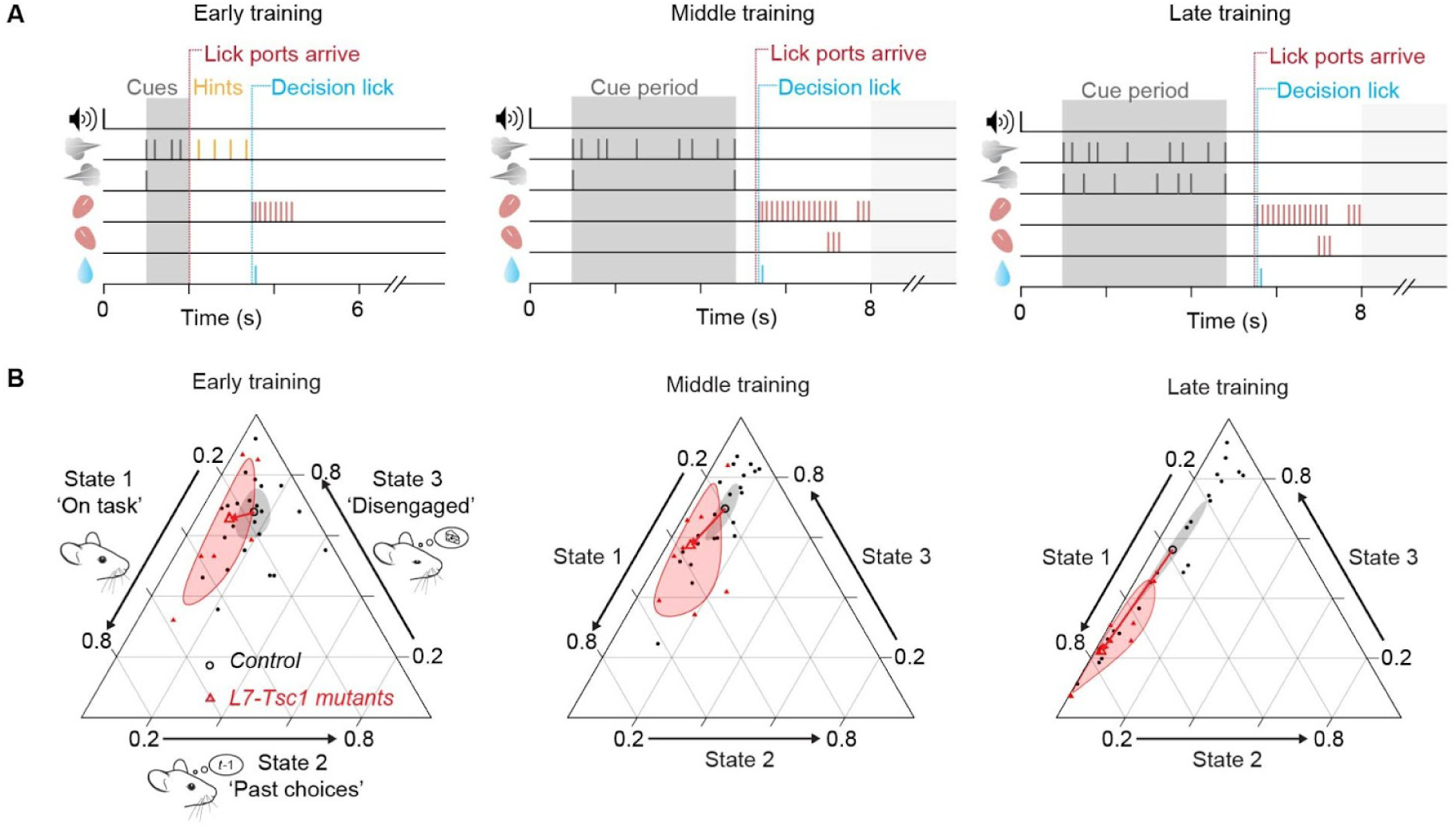
Cerebellar mutant mice show increasing on-task focus as training progresses (A) Three different training stages of the evidence-accumulation task. (B) Ternary plots for the composition of latent states with 95% confidence bands (shaded regions) around the compositional mean (open symbols) for *L7-Tsc1* mutant mice and control mice. Each data point represents one mouse. The 95% confidence regions are based on an assumption of normality for compositions. Arrows indicate the shift from the average of the control group to the average of the *L7-Tsc1* mutant mice.

**Figure 5.**
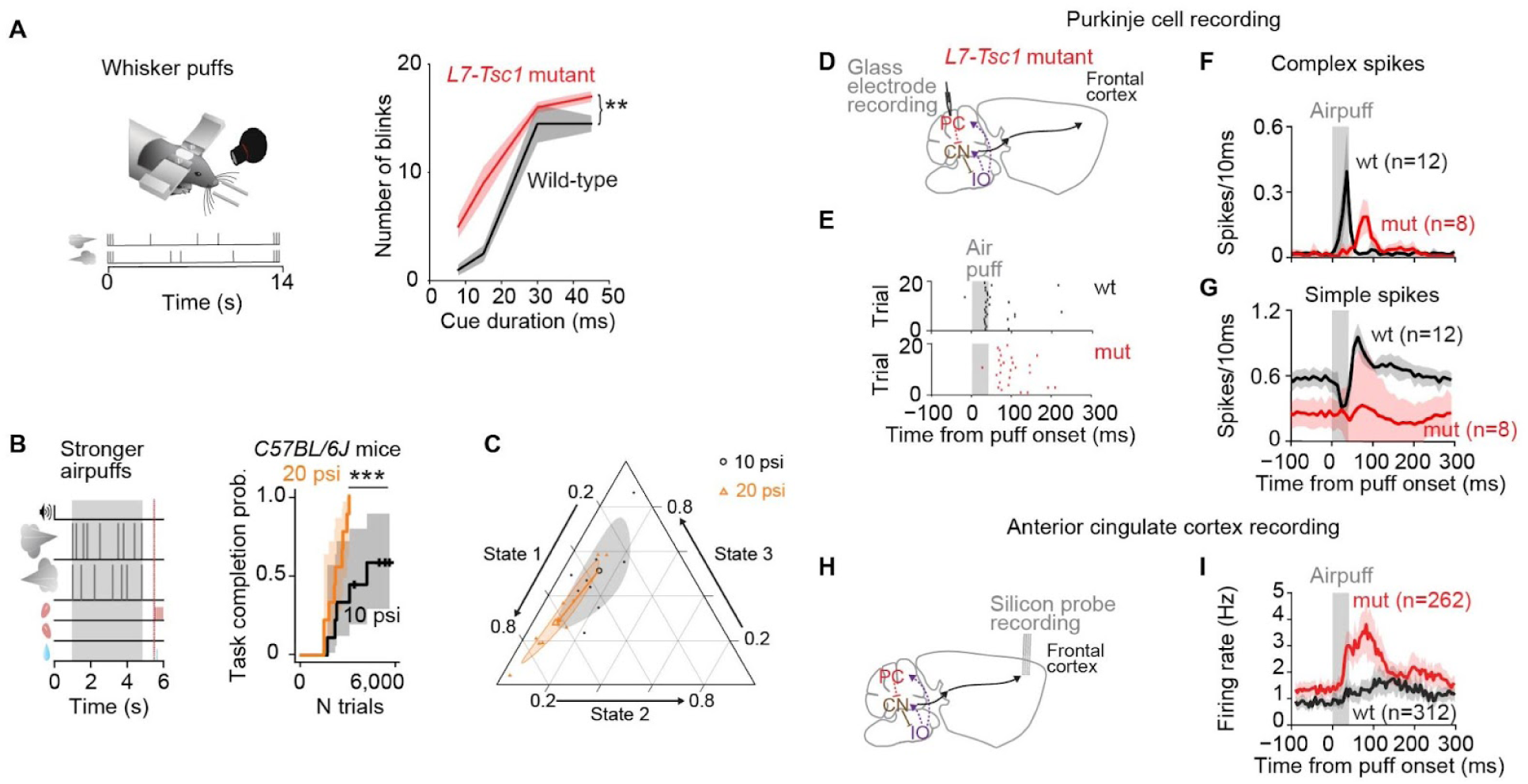
Prolonged whisker puff responses in awake behaving *L7-Tsc1* mice in cerebellar complex spikes and forebrain (A) Left: sensory sensitivity test with bilateral and unilateral whisker puffs in wild-type C57BL/6J mice. Right: median number of blinks in response to puffs of different durations for *L7-Tsc1* mutant mice (*n* = 16) and wild-type littermates (*n* = 7). Shaded areas indicate the estimated s.e.m. (B) Increased salience through stronger puffs leads to enhanced learning. Kaplan-Meier estimator of task completion for animals receiving standard (10 psi, 9 mice) or stronger (20 psi, 9 mice) puffs. Shaded areas represent 95% confidence intervals. (C) Ternary compositional plot of latent states with 95% confidence bands (shaded regions) around the compositional mean (open symbols) at middle levels of training. The arrow indicates the shift from the 10 psi average to the group receiving stronger airpuffs. (D) Schematic of recording site for whisker puff responses in *L7-Tsc1* mutant mice. (E) Example raster plots of Purkinje cell complex spikes during 20 trials from one wild-type animal (top) and one *L7-Tsc1* mutant animal (bottom). (F-G) Average number of spikes per 10 ms bins in response to an airpuff to the whiskers (data from 12 cells in 4 *L7-Tsc1* mutants and 8 cells in 5 wild-type mice) for Purkinje cell complex spikes and simple spikes. (F) Delayed time course of the airpuff evoked response of complex spikes compared with wild-type littermates. (G) Delayed and smaller simple-spike response in mutant mice compared with wild-type littermates. (H) Schematic of silicon probe recording site in forebrain. (I) Average firing rates in anterior cingulate cortex in response to an airpuff to the whiskers (4 mutants and 5 wild-types). Shaded areas represent 95% confidence intervals.

### Increased sensory intensity is sufficient to accelerate learning

Because simple associative conditioning was not improved in *L7-Tsc1* mice (Figure 1D) and because evidence-accumulation learning trended faster than wild-type at multiple stages of training (Figure S1), we hypothesized that the accelerated learning rate might be related to processing steps other than the direct response to airpuff stimuli. In several mouse models, a variety of motor and nonmotor phenotypes have been shown to be regulated by enhanced sensory salience (Orefice et al., 2019), raising the possibility that enhanced saliency of sensory events might lead to accelerated learning as well in our task. We therefore measured sensory reactivity before training. Naive *L7-Tsc1* mutant mice showed enhanced blink responses to individual airpuffs (Figure 5A), as well as to auditory stimuli (Figure S5A,B), indicative of altered sensory processing. In the DTSC task, responses to unconditioned-stimulus nose taps in mutants also tended to be longer in duration (Figure S5C-F). Therefore *L7-Tsc1* mutant mice are hypersensitive across at least two sensory modalities.

To test whether increased sensory intensity would be sufficient to accelerate learning, we increased the intensity of airpuffs from 10 psi to 20 psi in wild-type C57BL/6J mice. This was sufficient to accelerate training to a degree similar to that seen in *L7-Tsc1* mutant mice (Figure 5B), and showed a tendency (*p* = 0.10) toward increased state 1 occupancy during middle stages of training (Figure 5C). The shape of the psychometric curves in trained mice was similar whether animals received 10 psi or 20 psi airpuffs (Figure S6A,B). This result suggests that naturally occurring sensory reactivity is sufficient to modulate processes somewhere in the brain (whether cerebellar or extracerebellar) to accelerate learning of an evidence-accumulation task requiring integration of sensory evidence.

### Mutant mice show enhanced response to task stimuli in crus I and anterior cingulate cortex

In response to puffs of different durations (Figure 5A), *L7-Tsc1* mutant mice (*n* = 16) blinked more times than wild-type littermates (*n* = 7; F_(1)_ = 7.44, *p* = 0.008; no interaction with puff-dependence (two-way ANOVA, F = 0.98, *p* = 0.4). Stronger puffs also led to faster learning (Figure 5B): 20 psi puffs (9 mice, median 2225 trials) led to completion of training in fewer trials than standard 10 psi puffs (9 mice, median 4275 trials; *χ*^2^_(1)_ = 7.11 *p* = 0.00047, log-rank test).

Stronger puffs also shifted behavior during learning toward the on-task state. The log ratio state 2/state 1 increased with training level (two-way ANOVA, F_(1)_ = 44.2, *p* < 0.001), and trended towards greater values at 20 psi (F_(1)_ = 3.5, *p* = 0.07). Post hoc comparisons using the Tukey HSD test indicated a trend towards significance for middle stages of training (levels 3-4, the earliest levels at which the manipulation occurred, *p* = 0.10; Figure 5C), but not for later stages of training (levels 5-6, *p* = 0.99). The log ratio state 3/state 1 was not different for either group (F_(1)_ = 1.4, *p* = 0.24) or training level (F_(1)_ = 0.8, *p* = 0.37). In summary, more intense puff stimulation makes mice blink more, stay on task, and learn faster.

To measure neural signals accompanying enhanced sensory responsiveness, we performed *in vivo* electrophysiological recordings from airpuff-responsive Purkinje cells in crus I (Figure 5D). In wild-type mice, sensory cues triggered complex spikes (Figure 5E,F), with a delayed simple-spike response (Figure 5G). To test whether cerebellum-to-neocortex pathways were enhanced in *L7-Tsc1* mice, despite the fragility of *Tsc1^-/-^* Purkinje cells (Tsai et al., 2012) we were also able to make behavioral and electrophysiological recordings of Purkinje cells in naive *L7-Tsc1* mutant mice (Figure 5D-I). In *L7-Tsc1* mutant mice, complex spikes were activated after a latency of 53±7 ms (mean±SD), longer than the 13±7 ms seen in wild-type littermates (*t*_(19)_ = −10.7, *p* < 0.0001). The time course of responses was delayed: the mean area under the curve (AUC) was higher for mutants during the 50-100 ms following airpuff onset (relative to baseline rate pre-stimulus: mutant mice AUC = 5.1±2.0, wild-type littermates AUC = 0.8±0.7, *t*_(19)_ = −6.3, *p* < 0.0001), while the AUC in the 0-50 ms interval was higher for wild-type mice (mutant mice AUC = 1.1±0.5, wild-type AUC = 7.1±2.0, *t*_(19)_ = 7.89, *p* < 0.0001). Thus *L7-Tsc1* mice share with optogenetically-stimulated mice a delayed increase in complex spiking.

Simple spikes (Figure 5G) also had a later onset (latency in wild-type mice 11±8 ms (mean±SD); mutant mice 29±15 ms, *t*_(19)_ = −3.3, *p* = 0.004). However, unlike complex spikes, simple-spike responses were reduced (wild-type mean AUC = 191±27, mutant mean AUC = 71±85, *t*_(19)_ = 4.3, *p* = 0.0004). As expected, this decreased simple-spike response also led to disinhibition of negative feedback from deep nuclei (Figure S7B) onto inferior olivary neurons (Kim et al., 2020; Llinás, 2014).

Influences of cerebellum are conveyed via long-range pathways that project throughout thalamus and neocortex (Pisano et al., 2021), including two associative regions implicated in decision-making (Chabrol et al., 2019; Gao et al., 2018; Kennerley et al., 2006) that receive substantial disynaptic input from crus I (Pisano et al., 2021): anterior cingulate (Figure 5H, I) and anterolateral motor cortex (Figure S7C). When we recorded from ACC in wild-type and in mutant *L7-Tsc1* mice, we found an increased AUC in mutant mice (relative to baseline rate pre-stimulus: 678±1109) compared to wild-type mice (398±646, *t*_(573)_ =-3.6, *p* = 0.0003). Thus in *L7-Tsc1* mutant mice, we find enhanced coupling of crus I to ACC. Secondly, we observed an enhancement in cue-evoked activity with a similar time course as the changes in complex-spike activity. In untrained wild-type mice, airpuff-evoked complex spikes reached 90% of the maximum firing rate at 25±8 ms (latency, 13±7 ms) followed by anterior cingulate at 137±83 ms (latency, 72±70 ms), which were significantly different from each other (peak firing: *t*_(271)_ =-4.5, *p* = 0.00001, latency: *t*_(271)_ =-2.8, *p* = 0.006). In mutant mice the order more closely spaced, with complex spikes reaching 90% of maximum firing rate at 73±5 ms (latency, 53±7 ms) followed by anterior cingulate at 106±78 ms (latency, 46±49 ms; peak timing: *t*_(316)_ =-1.1, *p* = 0.26, latency: *t*_(316)_ = 0.35, *p* = 0.73). This altered timing was not seen in the barrel field of the primary somatosensory cortex (Figure S7D). These results suggest that cerebellar complex spike output can influence neocortical activity and is thus available as a driver of learning.

### Cue-locked optogenetic stimulation also boosts learning rate

To determine whether perturbation of the cerebellum was sufficient to enhance learning, we directly manipulated neural activity in wild-type mice by expressing the optogenetic probe channelrhodopsin-2 in Purkinje cells (Figure 6). We optogenetically (Titley et al., 2019) reinforced each cue, starting after mice had passed out of the early stage of training, by pairing each sensory stimulus with a cue-locked ipsilateral optogenetic light flash applied over crus I (Figure 6A). Cue-locked ipsilateral flashes led to faster learning than in strain-matched controls not expressing ChR2 (Figure 6A; median 2512 trials in 6 mice, compared with 3311 trials in 5 wild-type littermates; *χ*^2^_(10)_ = 8.18, *p* = 0.0042, log-rank test). Analysis of variance on the log ratio state 2/state 1 yielded a significant difference for both group (F_(1)_ = 12.3, *p* = 0.003) and training level (F_(1)_ = 16.4, *p* < 0.001). Post hoc comparisons using the Tukey HSD test detected no difference in log ratio state 2/state 1 at middle stages of training (levels 3-4, the earliest levels at which the manipulation occurred, *p* = 0.43), and an increase by late stages of training (levels 5-6, *p* = 0.02). By late training, 4 out of 5 optogenetically-reinforced mice spent more than 90% of the trials in state 1 and fewer than 5% of the trials in state 2 (Figure 6B). The log ratio state 3/state 1 was lower in the optogenetically-reinforced mice than in animals not expressing ChR2 (F_(1)_ = 10.3, *p* = 0.006), without a difference by level, F_(1)_ = 0.09, *p* = 0.77). In trained mice, the shape of the psychometric curves in each state was similar with or without cue-locked optogenetic stimulation (Figure S4C,D). In summary, cue-locked optogenetic stimulation led to a gradual enhancement in the occupancy of state 1, which grew with training.

**Figure 6.**
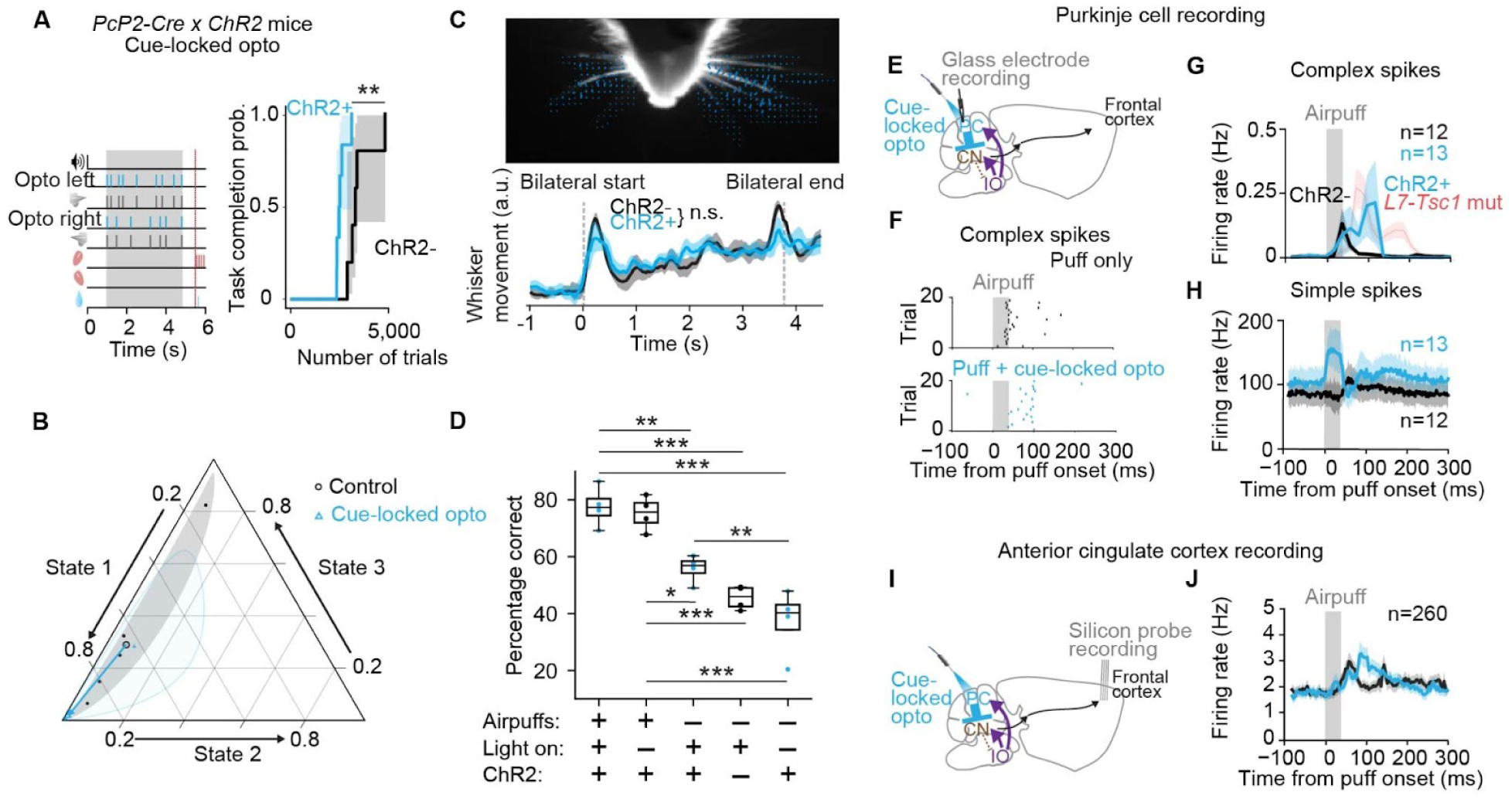
Altered whisker puff responses and faster learning with cue-locked optogenetic stimulation of Purkinje cells (A) The evidence-accumulation task and Kaplan-Meier estimator of task completion for 6 *Pcp2-Cre* × *ChR2* mice with cue-locked bilateral optogenetic activation of crus I. (B) Ternary compositional plot for the latent states with 95% confidence bands (shaded regions) around the mean (open symbols) at later levels of training. The arrow indicates the shift from the control average to the group receiving cue-locked optogenetic stimulation. (C) Detection of whisker movement during the evidence-accumulation task. Blue arrows indicate detection of whisker movement measured using a region-of-interest optical flow analysis. Shaded areas in the bottom plot include 95% confidence intervals. (D) Performance in the evidence-accumulation task in 4 trained *Pcp2-Cre* × *ChR2* mice for trials at level 7 under various experimental conditions, from left to right: airpuffs combined with cue-locked optogenetic activation of Purkinje cells in crus I, airpuffs only, optogenetic activation only, and negative controls lacking ChR2 or optogenetic stimuli. Due to anti-biasing parameters, chance-level performance for each condition was always less than 50%. (E) Cerebellar recording site in the whisker puffs + cue-locked optogenetic experiment. (F) Example raster plots of Purkinje cell complex spikes during 20 trials with only a whisker puff (top) or with a whisker puff paired with optogenetic stimulation (bottom). (G) Purkinje-cell complex-spike recordings (average data from 4 mice) show that pairing airpuffs with optogenetic stimuli led to a shift in the time course of activity toward later times after airpuff onset. (H) Simple spikes in the same Purkinje cells led to an increased firing rate without time shift. (I) Silicon-probe recording in forebrain combined with cue-locked optogenetic activation of cerebellar Purkinje cells. (J) Average firing rates in anterior cingulate cortex. The addition of optogenetic stimulation resulted in time-shifted responses to airpuffs. The red line in G indicates for comparison the time course of complex spike firing for *L7-Tsc1* mutant mice. Shaded areas represent 95% confidence intervals.

Optogenetically-driven acceleration of learning might arise from enhancement of direct sensory responsiveness, or alternately add an additional signal that shapes learning. In a sensory reactivity test, cue-locked optogenetic stimulation did not increase the number of blinks produced to higher-intensity airpuffs (within-mouse comparison; 25 psi puffs of 8-45 ms duration, F_(2,48)_ = 0.23, *p* = 0.80 for cue-locked flashes vs. no flashes, linear mixed-effect model, 5 mice). The amount of per-puff whisker movement in the evidence-accumulation task was also not affected (Figure 6C, rise of total frame-on-frame changed pixels during the cue period: *p* = 0.27, decay: *p* = 0.53, two-tailed Welch’s *t*-test, *n* = 5 ChR- mice and *n* = 6 ChR+ mice), indicating that the accelerated learning which occurs with cue-locked optogenetic stimulation drives mechanisms that are in some way separate from immediate sensorimotor processing.

We found that after training under cue-locked optogenetic enhancement to the highest level of task, mice were able to do the task both with airpuffs+opto (77.6% correct) and with airpuffs only (75.3% correct; Figure 6D). We were interested to observe that they could also perform above chance with optogenetic stimuli only (55.8%). Opto-only performance was, in turn, better than mice not expressing ChR2 (45.6% correct, *p* = 0.046, Conover post hoc test) or than mice not receiving optogenetic stimuli (37.2% correct, *p* = 0.0044). Together, these observations show that in well-trained mice, cerebellar activity alone is sufficient to play an autonomous role in evidence accumulation, but that under such conditions, cerebellar activity does not enhance the role of sensory evidence.

### Accelerators of learning converge on increased complex-spike and neocortical response

To measure neurophysiological consequences of cerebellar optogenetic stimulation, we made *in vivo* recordings in awake behaving naive ChR2-expressing mice (Figure 6E,I, Figure S7A). In these recordings, pairing airpuffs with cue-locked optogenetic stimulation led to an enhancement of simple-spike firing (Figure 6H) and a delayed increase in complex spike firing (Figure 6F,G). The enhanced complex-spike firing coincided with the end of the optogenetic stimulus, consistent with a disinhibitory effect of simple-spike firing on nucleo-olivary paths (Bengtsson and Hesslow, 2006).

Combining airpuffs with optogenetic stimulation resulted in a net delay in the time course of complex spikes compared with airpuffs-only (Figure 6G), with a smaller response during the first 50 ms following puff onset (airpuff+opto mean AUC = 0.4±0.3, airpuffs-only = 1.0±0.8; *t*_(24)_ = 2.5, *p* = 0.02, two-sided Student’s t-test) and a larger response in the next 50 ms (airpuff+opto = 1.8±0.3, airpuffs-only = 0.7±0.4; *t*_(24)_ = −7.5, *p* < 0.0001) and an accompanying decrease in the deep nuclear response (airpuff + opto: 10±6, airpuffs-only: 17±7, *t*_(17)_ = 2.3, *p* = 0.03; Figure S9A). Simple spikes were affected differently (Figure 6H): in the first 50 ms following airpuff onset, more simple spikes fired under the airpuff+opto condition (airpuff+opto mean AUC = 26±9, airpuffs-only = 14±5; *t*_(24)_ = −3.7, *p* = 0.001), but there was little measurable difference in the next 50 ms (airpuff+opto = 14±5, airpuffs-only = 16±5, *t*_(24)_ = 1.18, *p* = 0.25).

The use of wild-type mice allowed us to make silicon probe recordings in neocortex under cerebellar cue-locked optogenetic activation (Figure 6I, 3 mice). As was the case for the complex spike response, the addition of cue-locked optogenetic stimulation resulted in a delayed response in anterior cingulate cortex (Figure 6J), with a reduced response in the 50 ms following airpuff onset (relative to baseline rate pre-stimulus: airpuff+opto mean AUC = 124±212, airpuffs-only = 167±338, *t*_(259)_ = 4.6, *p* < 0.0001, paired t-test), and a larger response in the next 50 ms (airpuffs+opto = 200±390, airpuffs-only = 151±323; *t*_(259)_ = −4.3, *p* < 0.0001, paired t-test). We also observed an enhancement in anterolateral motor region (Figure S9B) activity, but not in the barrel field of the primary somatosensory cortex (Figure S9C). Thus cerebellar cue-locked optogenetic stimulation led to parallel changes in the time course of complex-spike cerebellar and associative-neocortical activity.

### Enhancements of learning are not correlated with simple-spike perturbations

So far we have shown that learning can be augmented by enhancing Purkinje cell activity during sensory stimulation, either via *Tsc1* knockout or by cue-locked optogenetic stimulation. Both of these perturbations generate similar post-stimulus increases in complex spike timing and neocortical activity. However, they also affect simple-spike firing, which also has effects outside the cerebellum.

Continuous optogenetic stimulation of Purkinje cells leads to specific increases in simple-spike activity, and such stimulation in crus I in trained mice during the cue period and delay period including the first lick (Figure S8A) is known to impair performance through forgetting of accumulated sensory evidence (Deverett et al., 2019). This perturbation did not affect the amount of per-puff whisker movement in the evidence-accumulation task (Figure S8D, rise: *p* = 0.61, decay: *p* = 0.33, two-tailed Welch’s *t*-test, *n* = 5 ChR- mice and *n* = 6 ChR+ mice). Even though such continuous stimulation increased overall simple-spike activity, it did not enhance simple-spike responses to individual sensory cues, and there was no change in complex spike firing (Figure S8E,F) or deep nuclear firing (Figure S8H) in response to whisker puffs and no detectable effect on the learning rate (Figure S8B) or whisker puff responses in forebrain areas (Figure S8G,I,J). There was also no change in state 1 occupancy, although there was a reduction in state 2 occupancy that was visible starting at the earliest stages of perturbation (Figure S8C). Psychometric curves in trained mice were also not affected by continuous optogenetic activation (Figure S6C,D). Thus, an experimental perturbation that affects simple-spike but not complex-spike firing does not accelerate learning.

### During training, cerebellar activation enhances anterior cingulate responses to evidence

With some effort (Figure 7A, S10) it was possible to make acute Neuropixels recordings and deliver optogenetic stimuli as mice performed evidence accumulation. In wild-type mice undergoing level 3-5 training, we identified 165 units in anterior cingulate cortex that responded to bilateral whisker stimulation marking the start of the trial (Figure 7B), which we then used to probe cerebellum-forebrain interactions. These units also responded to unilateral evidence puffs presented on either side (Figure 7C), as well as to cerebellar optogenetic stimuli (with a 20-30 millisecond delay; Figure 7B, C).

**Figure 7.**
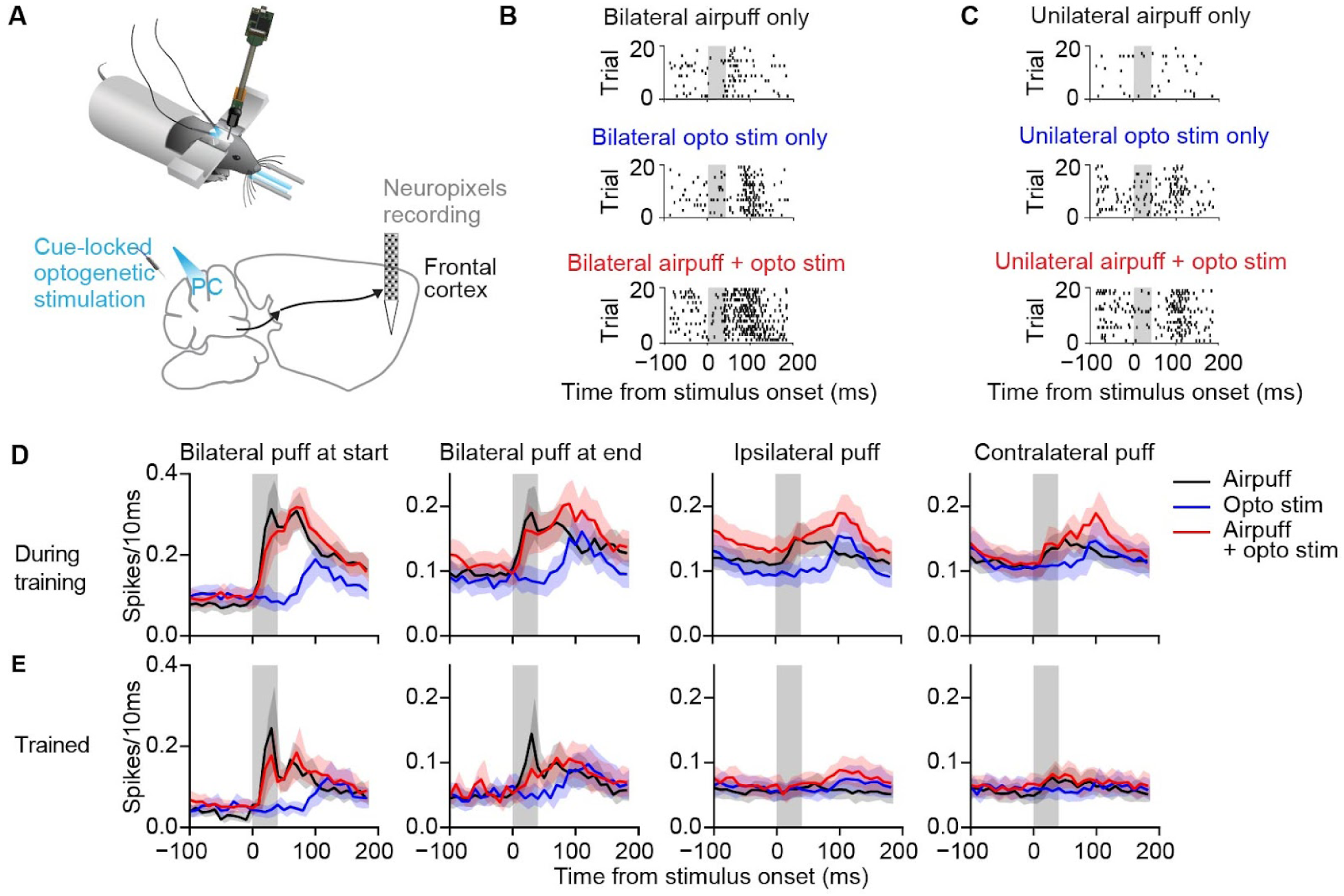
Neural responses in anterior cingulate cortex to optogenetic stimulation of the cerebellum and whisker puff stimuli during the evidence-accumulation task (A) Execution of the evidence-accumulation task during acute Neuropixels recordings and bilateral optogenetic stimulation of the cerebellum (top), and cerebellum and forebrain recording sites in addition to bilateral cue-locked optogenetic stimulation of Purkinje cells (bottom). (B) Spike rasters from an example single unit in anterior cingulate cortex during training in response to bilateral stimuli: airpuff only (top), optogenetic stimulation only (middle), or both (bottom). (C) Same as B, but for unilateral stimuli. (D) Average firing probability in response to a stimulus (black: airpuff only, blue: optogenetic stimulation only, red: airpuff combined with optogenetic stimulation), during levels 3-5 of training (*n* = 165 units from 3 recordings from 3 mice). (E) Same as D, but for highly-trained mice at level 7 or late stage of level 6 (*n* = 98 units from 3 recordings from 2 mice). Shaded areas represent 95% confidence intervals.

During a 200-millisecond post-stimulus period, pairing optogenetic stimuli with evidence airpuffs evoked a larger neural response than evidence airpuffs alone (Figure 7D; ipsilateral: 200-ms integrated response 27.8±24.7 for paired stimulation, 23.1±20.6 for airpuffs alone, *p* < 0.001; contralateral: integrated response 26.4±25.8 for paired stimulation, 22.8±20.4 for airpuffs alone, *p* < 0.001). This enhancement did not occur for the bilateral airpuffs that marked the start or the end of each trial. Thus cue-locked cerebellar optogenetic stimuli specifically enhance a forebrain effect of evidence airpuffs.

The completion of training (level 7 and late-stage level 6, *n* = 98 units) was associated with smaller responses in anterior cingulate cortex units to both unilateral and bilateral airpuffs (Figure 7E; two-way ANOVA, main effect of training vs. completed training with puff/opto/puff+opto as the second variate: start-of-trial bilateral puffs, F_(1)_ = 54.3, *p <* 10^−12^; end-of-trial bilateral puffs, F_(1)_ = 32.3, *p* = 2.0×10^−8^, ipsilateral puffs, F_(1)_ = 48.9, *p* = 7.4×10^−12^, contralateral puffs: F_(1)_ = 50.8, *p* = 3.0×10^−12^; no interaction effects for any stimulus, *p* > 0.1). Taken together, these results demonstrate that during training, cerebellar stimulation enhances ACC response to evidence airpuffs, and that this enhancement diminishes as training saturates.

## Discussion

Our experiments support the idea that cerebellar complex-spike output can accelerate nonmotor learning through increased on-task focus and modulation of brainwide responsivity to task variables. These cerebellar influences are transmitted to neocortical structures via major paths through thalamus and other midbrain structures (Fujita et al., 2020; Pisano et al., 2021) to support delayed activation and enhanced forebrain responses.

In *L7-Tsc1* mutant mice, inhibition of the medial prefrontal cortex has previously been found to improve social deficits and repetitive behaviors (Kelly et al., 2020). Among the extensive neocortical targets of cerebellar projections is the parietal cortex, where, interestingly, silencing of activity was recently shown to improve performance in evidence accumulation by reducing reliance on past evidence (Akrami et al., 2018). In addition, transcranial direct current stimulation of the right lateral posterior cerebellum in humans improves performance on a sentence completion task, as well as altering activity in multiple neocortical regions (Rice et al., 2021). Finally, manipulation of a crus I projection to the dentate cerebellar nucleus affects dopamine release in striatum (Washburn et al., 2024), and reduced Purkinje-cell activity can lead to enhanced post-inhibitory increases in cerebellar output to regulate the strength and timing of action (Witter et al., 2013). These routes provide ways for the cerebellum to influence forebrain function.

The differing consequences of cerebellar perturbation for learning of evidence accumulation and delay tactile startle conditioning (DTSC) are likely to arise from differences in their brain-wide neuronal substrates. Evidence accumulation is a more complex task than DTSC, as evidenced by two points. First, mice take more trials to complete learning of evidence accumulation than DTSC (see Figure 1D vs 1E). Second, evidence accumulation and DTSC call upon complex sensory integration/decision making vs. simple sensory-motor conditioning, respectively, and are likely to call on associative vs. sensorimotor brain areas. The evidence-accumulation task is known to depend on brain-wide networks including cerebellar crus I (Deverett et al., 2018; Pinto et al., 2018), whereas DTSC is an associative motor learning task which is dependent on anterior dorsal cerebellar vermis (Yamada et al., 2019), similar to classical cerebellar-dependent associative learning tasks such as delay eyeblink conditioning. The involvement of brain-wide networks in the evidence-accumulation task can explain why cerebellar manipulations impair DTSC acquisition, yet can either improve or impair evidence accumulation acquisition depending on the type and timing of the cerebellar manipulation and its effect on the forebrain (Figures 1E, 6A, S8B, and Deverett et al., 2019).

In this work we demonstrated the ability of optogenetic stimulation to reinforce task-related learning mechanisms. We found that delivering optogenetic stimuli in a cue-locked manner had two effects. First, if given during the training period, cue-locked stimuli could accelerate learning. Second, after training using cue-locked optogenetic stimuli, the same optogenetic stimuli were sufficient on their own to bias the mouse’s lick choices in the direction of stimulation. These findings show that it is possible for cue-locked optogenetic stimulation to boost both the effects of sensory stimuli for both learning and the learned evidence accumulation process.

Our results are consistent with the idea that complex spikes can enhance the salience of sensory stimuli, and that this enhancement can be transmitted to frontal cortex, including anterior cingulate, using rebound-firing-based mechanisms in the deep nuclei. These signals are larger during the learning of a complex task than after learning, and can account for the accelerated learning in *L7-Tsc1* mice because those mice, while impaired in their Purkinje cell function, still produce complex spikes but with a delay after the stimulus. These signals decrease as the animal becomes skilled at the task. We base this explanation on our comparison of the consequences of *L7-Tsc1* knockout on thalamocortical activity with the effects of focal crus I-specific cue-locked optogenetic enhancement. We found that with each sensory stimulus, the two perturbations had opposite effects on Purkinje cell simple spike and nuclear activity. However, during a longer period following each stimulus, both models led to enhancements in complex spikes and nuclear activity, as well as enhancements in ACC/ALM firing. Strong inhibition of cerebellar nuclear neurons is known to lead to post-inhibitory rebound (Llinás and Mühlethaler, 1988; Tadayonnejad et al. 2010), especially in lateral nuclei (Schneider et al., 2013), which is transmitted to thalamus and inferior olive. Thus our cue-locked optogenetic perturbation may replicate the *L7-Tsc1* condition’s effect on post-stimulus thalamocortical activity using a rebound deep nuclear mechanism.

Our work complements recent findings about roles for crus I in driving cognitive processing. Complex spike activity in crus I can encode cue identity or perceptual choices, and this activity is coupled with anterior cingulate cortex under social but not non-social conditions (Hur et al., 2023), suggesting that ascending transmission of information is actively regulated.

Indeed, crus I can tag salient moments such as on-goal events and perceptual choices (González et al., 2024; Yiu et al., 2025). In a striking parallel from motor learning, complex spikes can instruct plasticity in primary somatosensory cortex (Silbaugh et al., 2025). Our results suggest that such an instructive mechanism may act via rebound firing in the deep nuclei to entrain forebrain mechanisms, including systems that support evidence accumulation and decision-making.

Additional mechanisms not shared between the two perturbations might also hypothetically account for our findings. For example, although recorded Purkinje cells in *L7-Tsc1* mice showed an increase in airpuff-evoked simple-spike responses, they also showed reductions in the number of Purkinje cells (Figure 1B). The reduced overall Purkinje cell drive in *L7-Tsc1* mutant mice could account for our observation that both baseline and evoked deep-nuclear output were increased. At a later stage of processing, the effects of increased deep-nuclear output are transmitted to forebrain via nucleothalamic synapses, which have recently been found to be strengthened in *L7-Tsc1* mice (Nishiyama et al., 2024). These loci provide additional mechanisms for cerebellar alterations in *L7-Tsc1* mice to cause hyperresponsiveness in forebrain.

### The cerebellum and global coherence

The convergence of impaired associative motor learning, increased sensory sensitivity, and accelerated task learning in *L7-Tsc1* mice (Figure 8) echoes traits found in autism spectrum disorder (ASD). ASD is associated with islands of enhanced function, including perceptual domains and technical and even artistic capacities (Happé and Frith, 2006; Mottron et al., 2006). According to the weak central coherence account of ASD, these enhanced capacities can be explained by a detail-focused cognitive style in which individual perceptual features are emphasized. In the global coherence account of autism spectrum disorder, the capacity to extract global form and meaning is displaced by superiority on local or detail-focused processing (Happé and Frith, 2006). Our findings with cerebellar perturbations demonstrate one aspect of such processing, sensory hypersensitivity (Mottron et al., 2006), and an association with accelerated capacity to learn a sensory-integration task. In mouse models, enhancement of peripheral sensory sensitivity leads to alterations in central brain circuitry including ACC, as well as ASD-like motor and nonmotor phenotypes (Orefice et al., 2019, Lee et al., 2024).

**Figure 8.**
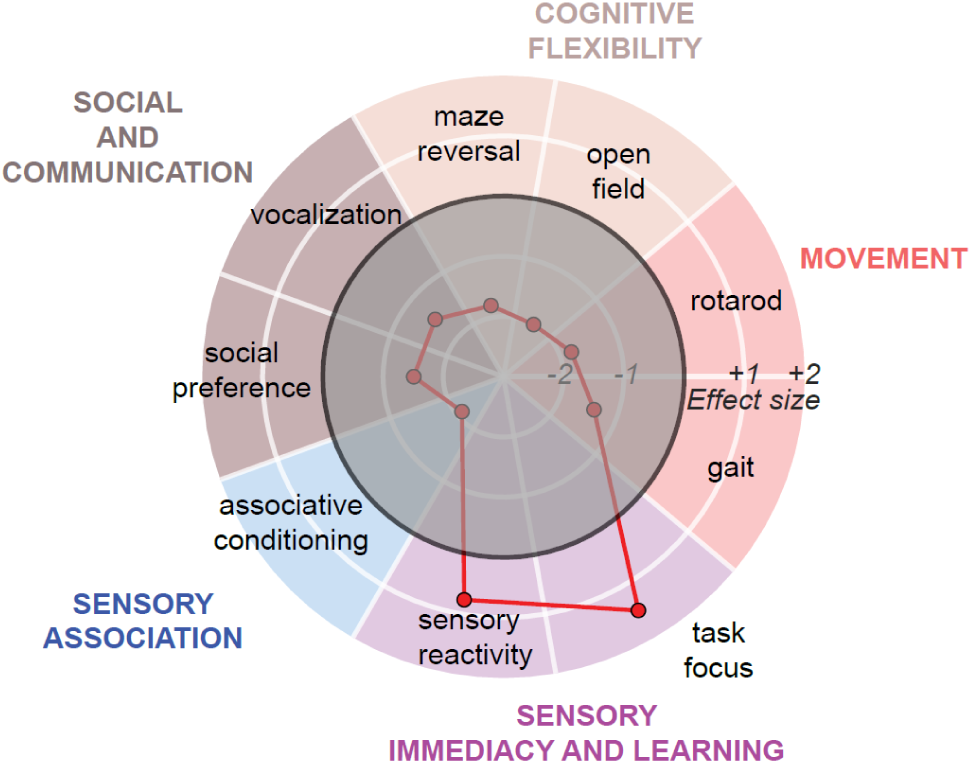
*L7-Tsc1* mutant mice express an island of enhanced sensory reactivity and task focus amid a variety of impairments Each dot represents an estimated effect size (Cohen’s *d*) for the behavior of *L7-Tsc1* mutant mice compared to their wild-type littermates. The thick circle indicates typical behavior (effect size 0). Based on data presented in this paper and from Tsai et al., 2012; Kloth et al., 2015; and Klibaite et al., 2021.

Our results show that cerebellar perturbation by itself can lead to one predicted feature of such increased sensitivity, hyperreactivity of neocortical circuits (Markram and Markram, 2010). In this way cerebellar disruption provides one neural substrate for a seemingly paradoxical phenomenon seen in ASD: broad disruption of cognitive and social function, accompanied by high performance in specific skill domains (Happé and Frith, 2006). This combination of features is reminiscent of atypicalities of sensation and perception reported in ASD, which have been interpreted in terms of a broadening of Bayesian priors about the sensory world. Such “hypo-priors” can account for a tendency among autistic persons “to perceive the world more accurately rather than [be] modulated by prior experience” (Pellicano and Burr, 2012). Expressed over development, the action of hypo-priors on forebrain circuitry could lead to lasting changes in regions that regulate social and cognitive function (Wang et al., 2014).

## Acknowledgements

We thank members of the Wang lab, the BRAIN CoGS consortium, and Michael Häusser for discussion and advice, G. Joseph Broussard, Caroline Jung, Junuk Lee, Laura Lynch, Dafina Pacuku, and John Wuethrich for help with experiments, Eva Peet for help with analysis, Jonathan Pillow for advice on latent-state analysis, Zahra Dhanerawala, Austin Hoag, and Sanjeev Janarthanan from the BRAIN CoGS Histology Core for brain clearing, imaging and probe localization, and Carlos Brody, Sabine Kastner, Manuel Schottdorf and Thomas Zhihao Luo for comments on the manuscript. The work was supported by National Institutes of Health grants R01 NS045193, R01 MH115750, and U19 NS104648 to S.W., and European Union’s Horizon 2020 research and innovation programme under the Marie Skłodowska-Curie grant agreement No 844318 to M.O.

## Author contributions

S.W., M.O., and M.K. conceptualized the research. M.O., S.W., M.K. and B.D. designed the experiments. M.O., M.K. and T.C. performed the experiments. M.O., M.K., and Y.C. did data analysis and visualization. S.J.V. provided the code for the GLM-HMM analysis. M.O. and S.W. did funding acquisition, project administration, and supervision. S.W. and M.O. wrote the original draft, and S.W., M.O., and M.K. reviewed and edited the paper.

## Methods

### Mice

Experimental procedures were approved by the Princeton University Institutional Animal Care and Use Committee (protocol 1943-19) and performed in accordance with the animal welfare guidelines of the National Institutes of Health and in line with the European Directive 2010/63/EU on the protection of animals used for experimental purposes.

Data came from 156 mice (males and females, 2-5 months of age at the start of experiments) of genotypes C57BL/6J (The Jackson Laboratory, Bar Harbor, ME, 40 animals), *Pcp2-Cre* for Purkinje-cell specificity and Ai27D for channelrhodopsin-2 (38 animals *Pcp2-Cre* × Ai27D, acquired from The Jackson Laboratory, stock #010536 (RRID:IMSR_JAX:010536) and #012567 (RRID:IMSR_JAX:012567), respectively), 3 *Pcp2-Cre* × Ai148 mice for Purkinje cell dendritic imaging (Ai148 line acquired from Hongkui Zeng, Allen Brain Institute), and *L7*^Cre^;*Tsc1^flox/flox^*mice (75 animals). To create these Purkinje cell specific *L7*^Cre^;*Tsc1^flox/flox^*mice, *Tsc1^flox/flox^* (*Tsc1^tm1Djk^*/J, The Jackson Laboratory stock #005680) mutant mice were crossed into L7-Cre mice (B6.129-Tg(Pcp2-cre)2Mpin/J, The Jackson Laboratory, stock #004146). Our control mice comprised all littermates arising from the breeding strategy with unaltered Tsc gene expression in the offspring. We included all such littermates: *Tsc1^flox/flox^* (9 mice for behavior, 9 mice for electrophysiology), *Tsc1^flox/+^* (11 mice for behavior, 8 mice for electrophysiology), *L7^Cre^;Tsc1^+/+^*(1 mouse for behavior, 1 mice for electrophysiology), and *Tsc1^+/+^*(2 mice for behavior, 1 mice for electrophysiology). Learning rates in the evidence accumulation task among these 4 control groups were not significantly different from one another (one-way ANOVA, F_(3)_ = 0.078, *p* = 0.97). Control behavior and neurophysiological measures were therefore reported as a single group for the *L7-Tsc1* mice. Because of differences in learning rates across genotypes (i.e. C57BL/6J, *L7-Tsc1*, and *Pcp2-Cre* × *Ai27D*), we made comparisons with suitably matched controls with conditions matched as closely as possible. Experimenters were blinded to the genotypes of the mice for the duration of the behavioral experiments.

All mice were group-housed in reverse light cycle to promote maximal performance during behavioral testing, which took time during the day. For long-term behavioral experiments, mice were housed in darkness in an enrichment box containing bedding, houses, wheels (Igloo and Fast-Trac; K3250/K3251; Bio-Serv; Flemington, NJ, USA), climbing chains, and play tubes during all experimental days. At other times, mice were housed in cages in the animal facility, in groups of 2-4 mice per cage. During experiments in which water intake was restricted, mice received 1.0-1.5 mL of filtered water per day plus half of a mini yogurt drop (F7577; Bio-Serv; Flemington, NJ, USA), and body weight and condition was monitored daily. Mice always had *ad libitum* access to food pellets.

### Surgical procedures

For all surgeries, mice were anesthetized with isoflurane (5% for induction, 1.0–2.5% for maintenance), and were given buprenorphine (0.1 mg/kg body weight) and rimadyl (5 mg/kg body weight) after surgery and were given at least 5 days of recovery in their home cages before the start of experiments, except for acute *in vivo* electrophysiology experiments when the animals were allowed to recover for at least two hours between the craniotomy and the acute recordings.

For mouse *in vivo* imaging experiments, a 3-mm-diameter craniotomy was drilled over the left posterior hemispheric cerebellum of *Pcp2-Cre* × Ai148 mice. In all imaged mice, a window composed of a cannula (Ziggy’s Tubes and Wires, 316 S/S Hypo Tube 9R GA. 0.1470/0.1490’ OD × 0.1150/0.1200’ ID × 0.0197’ long) glued (Norland Optical Adhesive 71) to a glass coverslip (Warner Instruments 64–0720) was cemented atop the craniotomy, and then a custom-machined titanium headplate was cemented to the skull using dental cement (C and B Metabond, Parkell Inc.).

For optogenetic experiments, a headplate was implanted as described above, after which two ∼500 μm diameter craniotomies were drilled over the cerebellum, one over each hemisphere, directly posterior to the lamboid suture and ∼3.6 mm lateral to the midline in either direction. During optical fiber placement we identified crus I starting first with coordinates (posterior 0.5 mm and lateral 3.6 mm from lambda, referenced relative to a diagonal approach perpendicular to the bone) and then adjusting aim based on the appearance of blood vessels on the surface of the brain. In initial experiments, location in crus I was confirmed histologically. Ferrule implants were constructed with 400-μm-diameter optical fiber (Thorlabs FT400EMT) glued to 1.25-mm OD stainless steel ferrules (Precision Fiber Products MM-FER2007-304-4500) using epoxy (Precision Fiber Products PFP 353ND). Ferrules were positioned over each craniotomy with the fiber tip at the surface of the dura mater, and Vetbond (3 M) was applied surrounding the exposed fiber. Dental cement was then applied to secure the ferrule to the skull. Implants were cleaned before each behavior session using a fiber optic cleaning kit (Thorlabs CKF).

For *in vivo* electrophysiology, a headplate was implanted as described above, and a 2 mm craniotomy was drilled over the area of interest and the dura removed. For recordings from neocortex, the following stereotaxic coordinates were used: anterior cingulate cortex: ML 0-0.5 mm, AP 0.5-1.5 mm, DV 0.7-1.0 mm, anterolateral motor cortex: ML 1.5 mm, AP 2.5 mm, DV 0.7-1.0 mm, and barrel field of the primary somatosensory cortex: ML 2.5 to 3.5 mm, AP −0.8 to −1.8 mm, DV 0.6 to 1.5 mm. Two stainless steel screws for ground and reference wires (000–120 1/16 SL bind machine screws, Antrin Miniature Specialties) were inserted in the skull above the forebrain as far away from the craniotomy as possible. For cerebellar recordings, a small hole was drilled for a reference electrode in the interparietal bone at the midline. Craniotomies (0.5 mm by 1.0-1.5 mm) were made next to the intersection of the interparietal and occipital bones and for silicon probe recordings made as lateral as possible (ML 2.8 mm) to reach crus I, and for glass electrode recordings centered over the left and right lobule V and simplex, allowing a diagonal antero-posterior approach to reach crus I (see Figure S7A). Craniotomies were covered with Kwik-Cast silicone adhesive (World Precision Instruments) until the time of the recording.

### Behavior experiments

Mice were trained to perform an evidence-accumulation decision-making task as described previously (Deverett et al. 2018; 2019). The behavioral apparatuses were controlled by custom-written Python software as published previously (Deverett et al. 2018) (https://github.com/wanglabprinceton/accumulating_puffs). Animals were trained for 1.5-9.0 weeks, 7 days/week.

Briefly, head-fixed mice were seated in a tube for daily 1 h behavioral sessions consisting of 200-300 trials. In each trial, independent streams of randomly timed 40 ms airpuffs of 10 psi (unless otherwise indicated) with a minimum 200 ms interpuff interval were delivered to the left and right sides over the course of a 1.0-3.8-second cue period. Each trial begins and ends with a bilateral puff to mark the cue period.

After a delay period of 200-800 ms, lick ports were advanced into the reach of the animal, and animals received a 4 µl water reward when they licked to the side with the greater number of puffs. The animal’s decision was interpreted as the side licked first, regardless of subsequent licks. Anti-biasing procedures (Deverett et al. 2018) result in chance levels being <50%. To increase motivation, restriction of water intake started at least 5 days before the start of training and continued throughout the whole training period. Error trials included a buzzer sound and a prolonged inter-trial interval.

Animals went through different levels of training (levels 0-6) to reach the final version of the task (level 7). The very first time the animal is in the setup, they need to learn that if they touch the lick spouts with their tongue, they will receive a drop of water. Therefore, in level 0 only, any first lick of each trial is rewarded with a drop of water, regardless which side the animal licked on or which side was correct for that trial. In levels 1 and 2, guide puffs occur at a regular interval (2.5 Hz) only on the correct side after the cue period ends and until the animal makes a first lick. Guide puffs are meant to be a hint to the animal as to where the correct side is, and allow the animal to base their decision on the most recent airpuff only (see Figure 4A left). Mice automatically proceeded to the next level once they reached pre-defined performance criteria (see Table S1 for details of each level as well as the performance criteria). The time it took an animal to learn the task was defined as the total number of trials to reach level 7. Psychometric curves were fitted with the psychofit module (Carandini and Wells, 2022).

Light for optogenetic stimulation during the evidence-accumulation task was delivered as described previously (Deverett et al. 2019). Cue-locked optogenetic cerebellar activation occurred unilaterally, at the same side and time at an airpuff, at an intensity of 1.5-5.0 mW for a duration of 40 ms (generated by Master-8, A.M.P.I.). Continuous optogenetic activation occurred bilaterally with 5-ms pulses at 50 Hz throughout the entire cue period, delay period, and ended upon first lick contact. When optogenetic activation was used to manipulate the learning rate, the optogenetic activation only started from level 3, and at every trial from then on. When optogenetic activation was used to manipulate performance in trained mice, light was on in 20% of trials. In this case, analysis compares light-off and light-on trials only from behavioral sessions in which light was delivered.

For the delay tactile startle conditioning (DTSC) task (Yamada et al., 2019; Broussard et al., 2022), mice learned to elicit a startle (backward) movement in response to an initially neutral conditioned stimulus (CS; 250 ms; 5 mm 395-400 nm UV Ultraviolet LED, EDGELEC) that was paired to co-terminate with a startle-eliciting unconditioned stimulus (US, 50 ms tactile stimulus on the nose by taping foam to the stepper motor shaft (High Torque Nema 17 Bipolar Stepper Motor 92oz.in/65Ncm 2.1A Extruder Motor, Stepper Online); CS-US inter-stimulus interval, 200 ms). Task performance was defined as the fraction of trials with a conditioned response (CR) and completion of task training was defined as the generation of a response of at least 20 steps on the rotary encoder on at least 60% of trials (Broussard et al., 2022).

For sensory sensitivity tests, naive animals were head-fixed in a similar setup to the evidence-accumulation setup and received either whisker puffs or auditory cues. Animals were not trained nor expected to do anything in response to the sensory cues, and did not receive any rewards throughout the session. Animals received cues in sequences of in total 24 cues starting and ending with three cues with 200 ms inter-cue interval, and in between those, cues at random intervals (ranging from 0.8 to 3.0 s). Animals first received a sequence with cue durations of 8 ms, followed by sequences with longer cue durations (15, 30, and 45 ms for whisker puffs, and 15, 30, 45, 90, 180, 320, and 640 ms for auditory cues). Animals either received bilateral airpuffs to the whiskers at 20-25 psi, or auditory cues at 12 kHz. During sensitivity tests with whisker puffs, white noise was on in the background throughout the experiment. To determine eye blink responses, movies of the right side of their face and body were acquired using two USB cameras (Playstation Eye), modified by removal of infrared filters and encasings. Images were acquired at 30 Hz with 320 × 240 pixel resolution. Illumination was provided by an infrared LED array (Yr.seasons 48-LED Illuminator Light CCTV 850 nm IR Infrared Night Vision). Airpuffs were produced by activation of solenoids (NResearch, standard two-way normally closed isolation valve, 161T011) with input from an air source (ControlAir Type 850 Miniature Air Pressure Regulator). Air was delivered via two tubes custom-machined with uniform openings, and positioned parallel to one another, parallel to the anteroposterior axis of the animal, 10 mm apart mediolaterally and ∼1 mm anterior to the nose of the animal. Auditory cues were delivered to the apparatus by a speaker (Sony Tweeter XS-H20S) mounted below the apparatus. Analysis of eye blinks was performed using FaceMap (https://github.com/MouseLand/facemap; Stringer et al., 2019) with manual curation and further analysis in Python.

### Two-photon imaging

The procedure for two-photon imaging was similar to Deverett et al., 2018. Briefly, imaging was done using a custom two-photon microscope (Sutter Instrument Company, movable objective microscope with resonant scanning) under separate computer control using the MATLAB software ScanImage 2015 (Pologruto et al., 2003) (Vidrio Technologies, RRID: SCR_014307). Excitation light was provided by a Mai Tai Sapphire laser (Spectra-Physics) at 920 nm. A 16x objective lens (Thorlabs, 16X Nikon CFI LWD Plan fluorite objective, 0.80 NA, 3.0 mm WD) was used with ultrasound gel (Sonigel, Mettler Electronics) as the immersion medium. Excitation power measured at the output of the objective lens ranged from 10 to 50 mW. Images were acquired at 28 Hz, 512 × 512 pixel image size.

*Synchronization.* Two forms of synchronization signal were sent from the behavior computer to the imaging computer during imaging. The first was a TCP/IP signal indicating animal and session identity. The second was an I2C-based signal routed through a National Instruments card (NI USB-8451). Signals were sent at multiple timepoints throughout each trial, delivering information corresponding to individual defined moments in the trial, which were then embedded in microscope image frames via ScanImage I2C functionality and retracted with imfinfo Matlab function.

*Imaging data processing.* Imaging data were preprocessed, motion corrected and analyzed with Python package - CaImAn (https://github.com/flatironinstitute/CaImAn). Motion corrected movies were used for manual regions of interest (ROI) selection. The manually selected dendritic ROI were subsequently refined using the Seed Constrained Nonnegative Matrix Factorization (CNMF) with external masks algorithm (Giovannucci et al., 2019). After all ROI were refined with CNMF, the mean activity of all pixels in the ROI was computed for each frame, yielding raw time series data. ΔF/F0 was then computed from these raw traces, with baseline F0 being computed as the minimum of a median-filtered (1 s kernel) 12 s sliding window preceding each time point. Dendritic data analysis included trials of all cue period durations.

*Linear model with mixed effects (LME)*. For LME analyses in Figures 3, we used the MATLAB function ‘fitlme’ to fit data to two linear mixed-effect models. We fitted the number of dendritic calcium transients C_cue_ during the cue periods in a single imaging session to an LME that incorporated the number of air-puffs in the trial P, the latent behavioral state S, and their interactions as fixed effects, with latent state 3 set as the reference. Mouse and dendrite labels were included as random effects, giving the following equation:

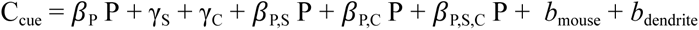

Where the β’s represent slope coefficients, the γ’s represent constants and *b*’s represent random effects. For each imaging session, we also modeled the Z-scored number of dendritic calcium transients during the 800 millisecond post-decision period following the lick (Z_decision_), using as inputs the outcome type Err (categorical: Correct or Error trial), latent behavioral state S, dendrite cluster C, and their interactions as fixed effects. The reference conditions were correct outcome, latent state 3, and the non-clustered dendrites. The resulting fit equation was

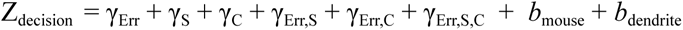

where these γ’s are different from those used in the C_cue_ model. Model outputs are available on our Github page.

### *In vivo* electrophysiology

For acute recordings from awake behaving mice, animals were head-fixed over a freely rotating cylindrical treadmill and the craniotomy site was opened by removing the Kwik-Cast plug and then filled with saline. Recordings were performed using either silicon probes for neocortex or glass electrodes for cerebellum, as described below. airpuffs to the whiskers were delivered by a pressure injector system (Toohey Spritzer, Toohey,

Fairfield, NJ, USA) which received signals from a signal generator (Master-8; AMPI) with an intensity of 20 psi and a frequency of 1 Hz, except for experiments with continuous optogenetic activation throughout the entire cue and delay period, when airpuffs were delivered with a frequency of 0.2 Hz. Mice received unilateral airpuffs ipsilaterally to the recording site for Purkinje cells, anterior cingulate cortex, and anterolateral motor cortex, and contralaterally to the recording site for cerebellar nuclei and the barrel field of the somatosensory cortex. For recordings with optogenetic stimulation, light onset started at the same time as the airpuff for the duration of the airpuff (40 ms) unless indicated otherwise. In a subset of experiments (Figure S9D) light started at the same time as the airpuff but remained on for longer (250 ms).

For neocortical recordings, a 64-channel silicon probe (Figures 1, 5, and 6; Neuronexus, A4×16-5mm-50-200-177 or A2×32-Poly5-10mm-20s-200-100) or a Neuropixels 1.0 probe (Figure 7) covered in Vybrant™ CM-DiI Cell-Labeling Solution (V22888; Invitrogen) was slowly placed above the craniotomy and lowered into the brain using a motorized micromanipulator (MP-225; Sutter Instrument Co.). The Neuronexus silicon probes were connected to two amplifier boards (RHD2132, Intan Technologies) using a dual headstage adapter (RHD2000, Intan Technologies). Recordings were made using an Open Ephys acquisition board at a sampling rate of 30 kHz. Neuropixels data were acquired using SpikeGLX (https://billkarsh.github.io/SpikeGLX/) and signals were digitized at 30 kHz. The probe was connected via a headstage to a PXIe data acquisition card (National Instruments, PXI-6133) in a PXI chassis (National Instruments, PXIe-1071), and connected to a computer via a PXIe interface (National Instruments, PXIe-8381). For all silicon probe recordings, high-pass filtering of the raw data at 300 Hz, common median referencing, and automatic spike sorting was achieved using Kilosort 2 (Pachitariu et al., 2016) (https://github.com/cortex-lab/Kilosort). Spikes were further manually curated using the Phy GUI (https://github.com/kwikteam/phy).

Single-unit recordings of Purkinje neurons and cerebellar nuclei neurons were performed using borosilicate glass electrodes (1B100F-4, World Precision Instruments) with 1- to 2-μm tips, short for Purkinje cells or very long gradual tapers for cerebellar nuclei cells, and 3 to 12 MΩ impedance, fabricated on a pipette puller (P-2000, Sutter Instruments Co.) and filled with sterile saline. The electrode was lowered into the cerebellum using an electrode holder that was positioned at a 40 or 90° angle to the craniotomy and controlled by a motorized micromanipulator (MP-225; Sutter Instrument Co.). The obtained electrical signals were amplified with a CV-7B headstage and Multiclamp 700B amplifier, digitized at 10 kHz with a Digidata 1440A and acquired in pClamp (Axon Instruments, Molecular Devices) in parallel with transistor-transistor logic (TTL) pulses from a signal generator (Master-8; AMPI) and with signal from pressure injector system (Toohey Spritzer, Toohey, Fairfield, NJ, USA).

Purkinje neurons were identified by the presence of complex spikes followed by a characteristic pause in simple spikes. For simple spike analysis, we required a minimum recording duration of 10 s, and the presence of at least one complex spike to confirm the identity of the Purkinje cell. For complex spike recordings the minimal recording duration was 100 s. In wild-type mice, recordings were more stable, resulting in equal numbers of recordings for simple spike and complex spike analysis. In mutant mice, recordings were less stable and reduced the relative number of analyzable complex spike recordings. We selected airpuff-responsive recordings for further analysis. The cerebellar nuclei contain a high density of neurons that are deeper than, and well separated from, cerebellar cortical layers, and showed clear single unit spike activity. Spike detection was performed using custom code written in MATLAB 2019a. Latency was calculated as the time where the change in firing rate reached 30% of the first peak (minimal value within 100 ms after puff onset for simple spikes, maximum value within 100 ms after puff onset for all other areas). The area under the curve was calculated as the area between the baseline pre-stimulus firing rate and the firing rate of each unit in number of spikes per bin, for the timespan indicated in the text.

### Histology

Animals were anesthetized with an overdose of ketamine (400 mg/kg)/xylazine (50 mg/kg) (i.p.) and transcardially perfused using a peristaltic pump with phosphate buffered saline (PBS) with 10 mg/ml heparin (Sigma H3149-100KU), followed by chilled 10% formalin (Fisher Scientific). Brains were extracted from the skull after perfusion, postfixed overnight at 4°C, washed and stored in PBS at room temperature. To visualize the probe locations using the CM-DiI track, brains were cleared and imaged by the BRAIN CoGS histology core facility. All brains underwent the same abbreviated iDISCO+ clearing protocol as previously described (Pisano et al. 2021). In short, after an overnight fix in 4% PFA, brains were rinsed in PBS at room temperature for four 30 minute sessions. Immediately brains were dehydrated 1 hr at each ascending concentration of methanol (20, 40, 60, 80, 100, 100%) and placed overnight in methanol at room temperature. The next day, they were being placed in 66% dichloromethane (DCM)/33% methanol for 3 hrs at room temperature. Brains were cleared with 100% DCM for two 15 min steps then placed in 100% benzyl ether (DBE). Brains were kept in fresh DBE prior to imaging and after for long-term storage. Tissue was imaged using a light-sheet microscope (Ultramicroscope II, LaVision Biotec., Bielefeld, Germany).

For quantification of Purkinje cells, Purkinje cells were stained with calbindin. Animals were transcardially perfused as described above, and after postfixation were stored in PBS at 4°C until sectioning. Whole brain sagittal sections were cut at 90 μm and collected in 0.1 M PBS. Sections were processed for immunohistology by washing with PBS and incubating for 1 hr at room temperature in a blocking buffer (10% normal goat serum, 0.5% Triton in PBS) prior to a 2-day incubation at 4°C in PBS buffer containing 2% NGS, 0.4% Triton and the rabbit anti-calbindin-D-28K primary antibody (C7354; Sigma-Aldrich St. Louis, MO, USA; 1:1000). Sections were subsequently washed in PBS, incubated for 2 hr at room temperature in the PBS buffer with goat anti-rabbit Alexa Fluor 488-conjugated secondary antibody (A-11008; Thermo Fisher Scientific, MA, USA, Invitrogen; 1:400), mounted on glass slides and covered with Vectashield. Images were acquired on the epifluorescent microscope Hamamatsu Nanozoomer. Using NDP.view2 Plus software, individual lobules were identified and Purkinje cells were assigned to lobules for counting.

### Corticosterone measurements

Animals were food deprived for 12-24 hrs before blood collection. Immediately after receiving airpuffs to whiskers at 20 - 25 psi in a headfixed setup for 10-20 minutes, ∼50 µl of blood was collected from the tail vein using a capillary tube, and then immediately disposed of in a heparin-coated 1.5 ml eppendorf tube. Samples were stored on wet ice for maximum 4 hours, after they were centrifuged for 10 mins at 4 °C at 3000 rpm. Of each sample 2-10 µl of plasma was collected, placed in new non-coated 1.5 ml eppendorf tubes and stored at −80 °C. For each animal, two duplicate samples of 1 µl each were used to determine plasma corticosterone levels using the Corticosterone ELISA Kit (K014; Arbor Assays, Ann Arbor, MI, USA) according to the manufacturer’s protocol. Plate reading was done using an Infinite 200Pro (Tecan Life Sciences, Morrisville, NC, USA) with i-control software. Results from both duplicates were averaged to get one final corticosterone measurement per animal.

### Generalized linear model-hidden Markov model

The generalized linear model-hidden Markov model (GLM-HMM) is an analytical approach used here to identify underlying hidden behavioral states that underlie the progression of a long series of behavioral trials. The model combines a set of Bernoulli GLMs with a hidden Markov model (Ashwood et al., 2021; Calhoun, Pillow, and Murthy, 2019; Bolkan et al., 2022). For each trial, an animal is modeled to have a latent state that governs its strategy to process information in order to make the binary choice of which side to lick. Each state corresponds to a specific GLM with a unique weight vector of input variables. At the beginning of a session, an initial state probability governs the likelihood of starting in a given state. Between trials, the transition matrix of HMM defines the probability to change from one state to another. The output of GLM-HMM in each trial is calculated as the probability of a Bernoulli response (i.e. the probability of a rightward lick) based on both the latent state of the current trial and the input variables. Delta cues (Δcues) is the number of airpuffs on the right side minus the number of airpuffs on the left side. Guide airpuffs (‘hints’) are included. Previous choice 1 is the animal’s choice on the previous trial. Previous choice 2 is the animal’s choice of the trial prior to the previous trial. Previous reward is the side of the reward on the previous trial. Bias is an offset constant in each state that represents the tendency to lick rightward independent of other input variables. The trials used to calculate the psychometric curve of a latent state are selected to have a posterior probability for that state larger than 0.8. The state occupancy of a certain state is calculated as the fraction of trials whose posterior state probabilities are greatest for that state. The model was trained with data from 22 wild-type mice used in a previous study (Deverett et al., 2018a); such an approach for constructing a model using out-of-sample training data is commonly employed in order to avoid overfitting. Data were fitted using an expectation-maximization algorithm with code adapted from https://github.com/Brody-Lab/venditto_glm-hmm.

### Drift-diffusion modeling

The drift-diffusion model (DDM) framework is an analytical approach used to identify how mice accumulate multiple pulses of evidence from the whole cue period to make decisions. Here the previously used discrete evidence model was employed (Deverett et al., 2019). Briefly, the DDM was developed based on Brunton et al. (2013), where an accumulator a(t) tracks evidence during each trial. Right-side stimuli correspond to positive deflections, and left-side stimuli correspond to negative deflections. The trial’s choice is determined by the sign of the accumulator. The model incorporates noise (σ ^2^) in the accumulator, single-stimulus Gaussian variability (σ^2^), and memory drift (λ). A negative λ results in leaky accumulation, while a positive λ causes instability. The model also includes bias and lapse parameters, with the latter accounting for random responses in a subset of trials. The model was fitted using PBupsModel Julia 0.6.3 package (https://github.com/misun6312/PBupsModel.jl). Parameter fitting utilized automatic differentiation to maximize model likelihood trial-by-trial and over time, with the Hessian matrix ensuring positive semidefiniteness for valid fits. Each fit used 1000 repetitions, randomly initializing parameters and omitting 20% of trials. Median parameters, standard deviation and 95% range were derived from these repetitions. Model accuracy was assessed using Bayesian Information Criterion through cross-validation, predicting held-out trial choices based on the best-fit parameters.To simulate the trial-wise trajectory of the accumulator for the latent states, best fit parameters of DDM (Supplementary Table 2) was used and calculation run the model on the same timing of the 5 left and 3 right discrete puffs with time steps of 15 ms (Figure 2I).

### Compositional data analysis

Traditional statistical methods that treat compositional data as independent and identically distributed can lead to misleading results because they ignore the inherent constraints imposed by compositional data; for instance, the fact that the sum of proportions always equals one. Compositional data analysis is a specialized framework for compositional data that directly addresses the proportional nature of the latent state data. Compositional analyses and ternary plots were conducted using the R package compositions (version 2.0-8, see van den Boogaart and Tolosana-Delgado, 2008). Zero or missing state occupancy values were imputed with the smallest positive normalized floating-point number. To ensure the total occupancy across the three states summed to one, this value was subtracted proportionally from the state with the largest occupancy. To graphically represent the dispersion structure of latent state data, centers were calculated using compositional means. Confidence regions were determined based on total variance and radius derived as 1.96 of the radial standard deviation on a log scale from a Fisher distribution with (D-1) and (N−D+1) degrees of freedom.

### Statistical analysis and presentation

Statistical tests used are indicated throughout the text. All further analysis was done with custom-written code in Python 3 using Spyder (https://www.spyder-ide.org/), and R version 4.3.3 (https://www.r-project.org/) using RStudio (https://www.rstudio.com/). For every figure, * = *p* ≤ 0.05, ** = *p* ≤ 0.01, *** = *p* ≤ 0.001. Boxes show median/interquartile range, and whiskers extend to the farthest data point that lies within 1.5 times the interquartile range from the end of the box. All figures show within-batch comparisons.

### Effect size calculations

For comparison with previous literature, differences between *L7-Tsc1* mice and wild-type controls were calculated as the difference between means divided by the standard deviation. Standard deviation was taken as the standard error multiplied by the square root of the number of measurements. Measures from previous literature were vocalization, social preference, rotarod, and maze reversal (Tsai et al., 2012), associative conditioning and gait (Kloth et al., 2015), and open field (Klibaite et al., 2021).

### Code and data availability

Code used for data acquisition is available at https://github.com/wanglabprinceton/accumulating_puffs. All data that support the findings of this study will be publicly available upon publication.

## Supplemental information

Figure S1 - S10, Supplementary Tables 1 - 2, and Supplementary Movies 1 - 2 are available for this paper.

**Correspondence and requests for materials** should be addressed to S.W. or M.O.

**Supplementary Movie 1.** Video from one wild-type animal and one *L7-Tsc1* animal (littermates) performing two trials of the evidence-accumulation task.

**Supplementary Movie 2.** Example of two-photon calcium imaging of Purkinje cell dendrites during performance of the evidence-accumulation task.

---

**Figure S1.**
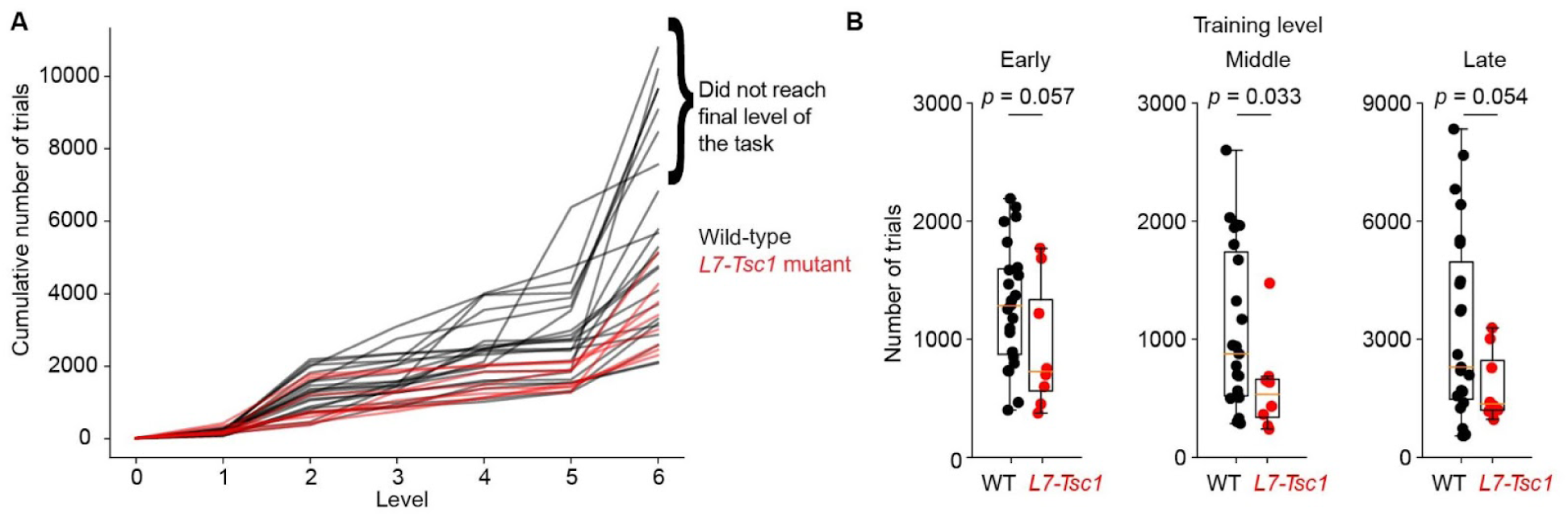
Learning trajectory for the evidence accumulation task for individual mice (A) Cumulative number of trials needed to graduate to the next level for *L7-Tsc1* mutant mice (red) and control mice (black) in the evidence accumulation task. Each line indicates an individual animal. Six control animals did not reach the final level of the task. (B) Number of trials needed at early training levels (0, 1, and 2), middle (3 and 4) and late (5 and 6) levels for *L7-Tsc1* mutant mice (red) and control mice (black) in the evidence accumulation task. Learning was faster at middle stages of training (levels 3 and 4, *t*_(30)_ = 1.9, *p* = 0.033, one-tailed t-test; Figure S1B) and also showed a tendency to be faster at early (levels 0, 1, and 2; *t*_(30)_ = 1.6, *p* = 0.057) and late (levels 5 and 6, *t*_(30)_ = 1.7, *p* = 0.054) stages.

**Figure S2:**
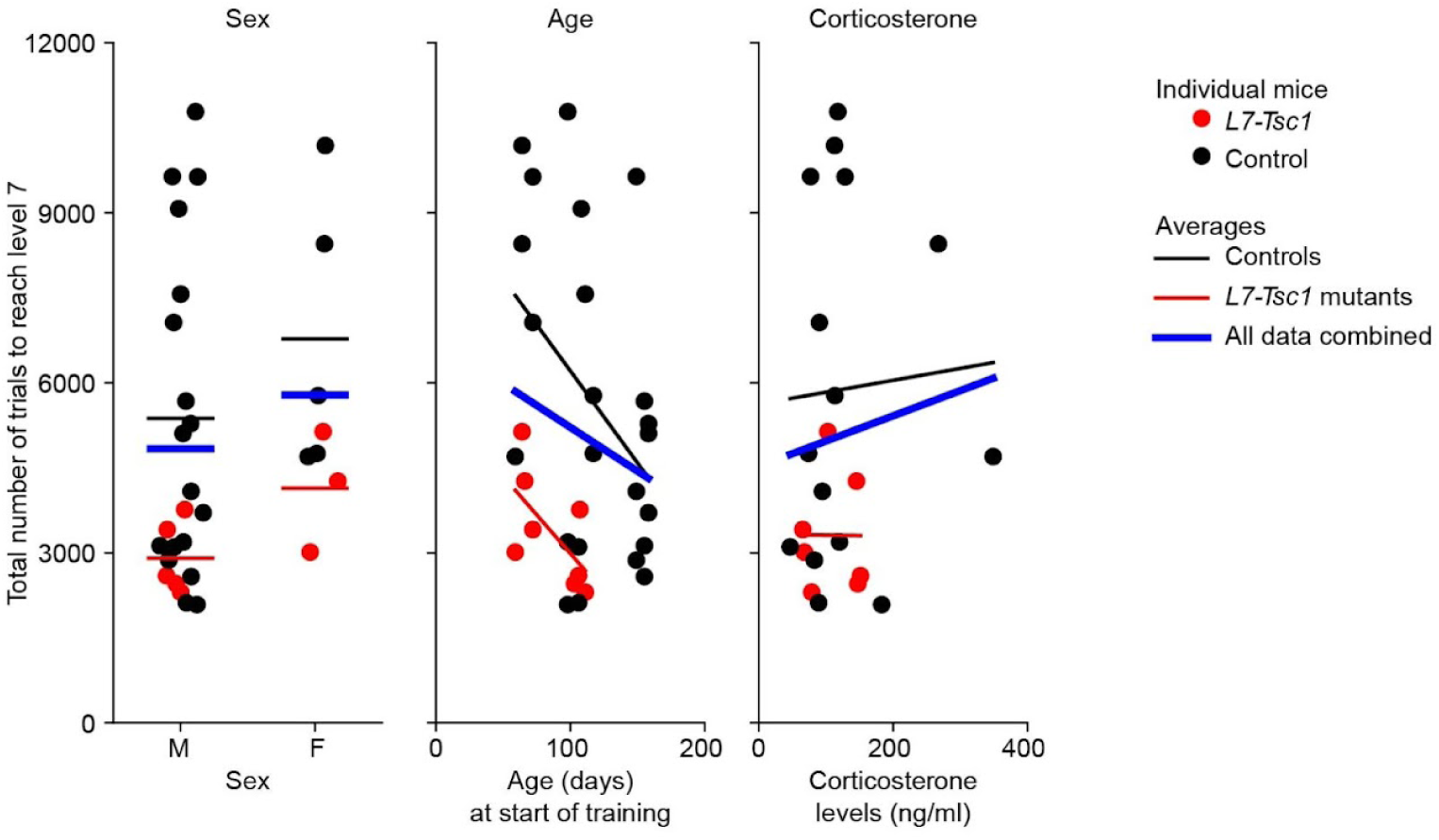
No correlation between the total number of trials to reach the final level of the evidence accumulation task and sex, age, or corticosterone levels. Numbers of animals are lower in the right panel than in the left and middle panels, as not all animals had blood samples taken to measure corticosterone levels.

**Figure S3.**
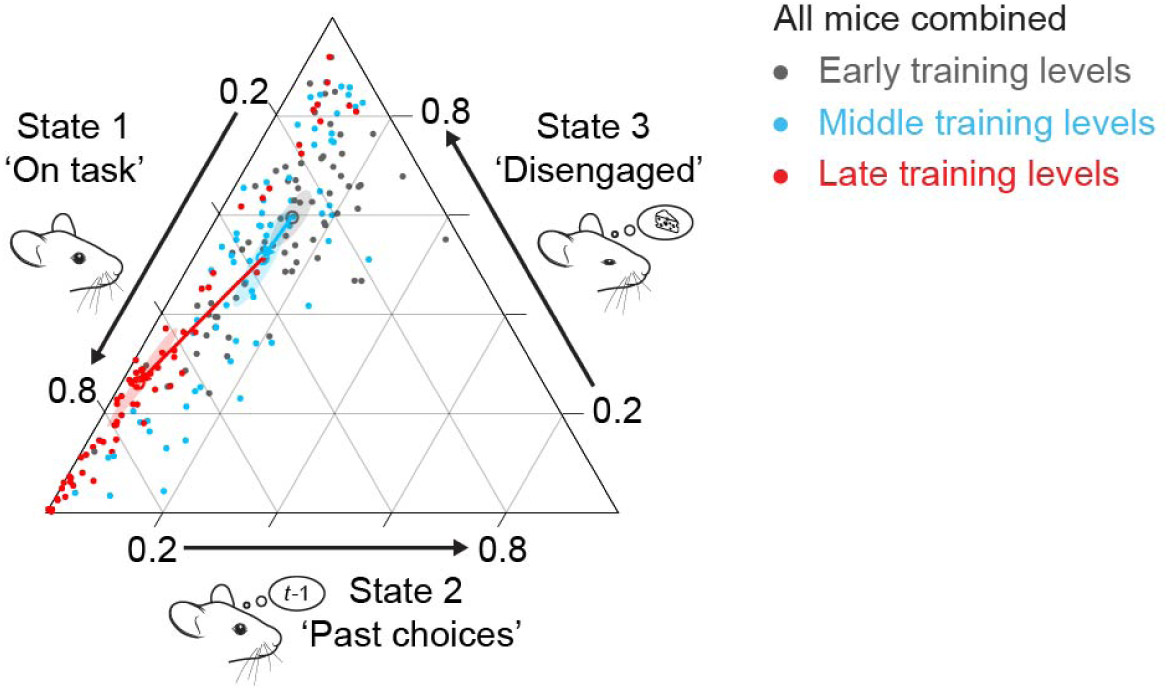
Altered state occupancy throughout training across all mice Ternary plot for the composition of latent states with 95% confidence bands (shaded regions) around the compositional mean (open symbols) for all mice combined (n=51, data pooled from Figure 4B, 5C, 6B, and S8C) across training levels (early: levels 0-2, middle: levels 3-4, late: levels 5-6). Analysis of variance on the log ratio state 2/state 1 yielded a significant effect for training level (F_(2)_ = 53.9, *p* < 0.001). Post hoc comparisons using the Tukey HSD test indicated no significant difference between early and middle stages of training (*p* = 0.30), but did show differences between early and late stages of training (*p* < 0.001), and between middle and late stages of training (*p* < 0.001). Analysis of variance on the log ratio state 3/state 1 also yielded a significant effect for training level (F_(2)_ = 5.3, *p* = 0.006). Post hoc comparisons using the Tukey HSD test indicated a significant difference between early and late stages of training (*p* = 0.004), but not between middle and late stages of training (*p* = 0.18), and early and middle stages of training (*p* = 0.30). Each data point represents one mouse. The 95% confidence regions are based on an assumption of normality for compositions. The arrows indicate the shift from the early to middle stages of training (blue arrow) and the shift from the middle to late stages of training (red arrow).

**Figure S4.**
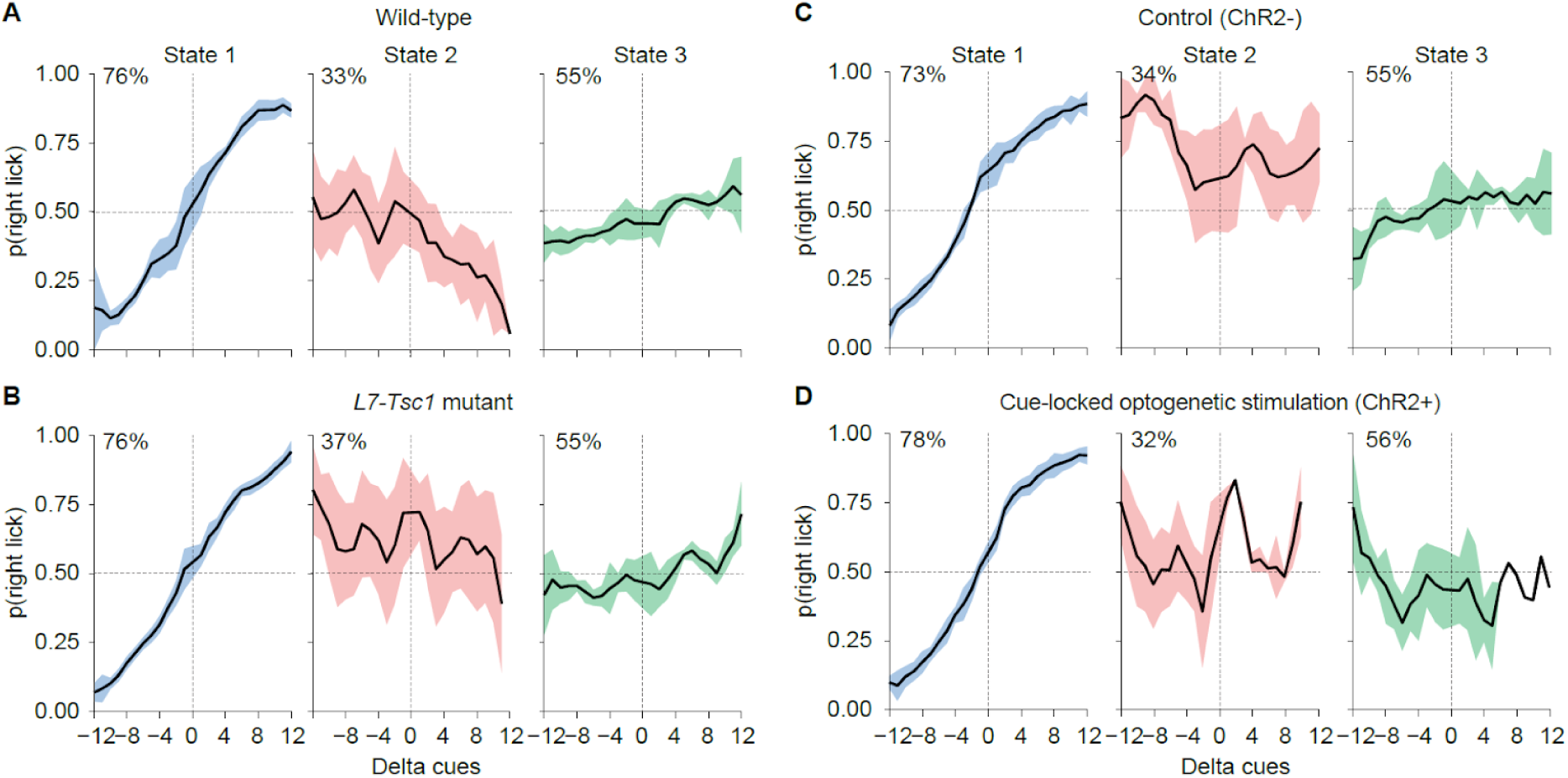
Psychometric curves in the three states remain the same for *L7-Tsc1* mice and for mice receiving cue-locked optogenetic stimulation (A-D) Psychometric curves for the three states, averaged across all wild-type mice (A), *L7-Tsc1* mutant mice (B), ChR- mice (C) and ChR+ mice receiving cue-locked optogenetic stimulation of Purkinje cells in crus I (D). In the top left of each plot is the percentage correct over all trials in that state. Missing data points or data points without error bars indicate none or one animal at that data point due to low state occupancy. Shaded areas represent 1 s.d.

**Figure S5.**
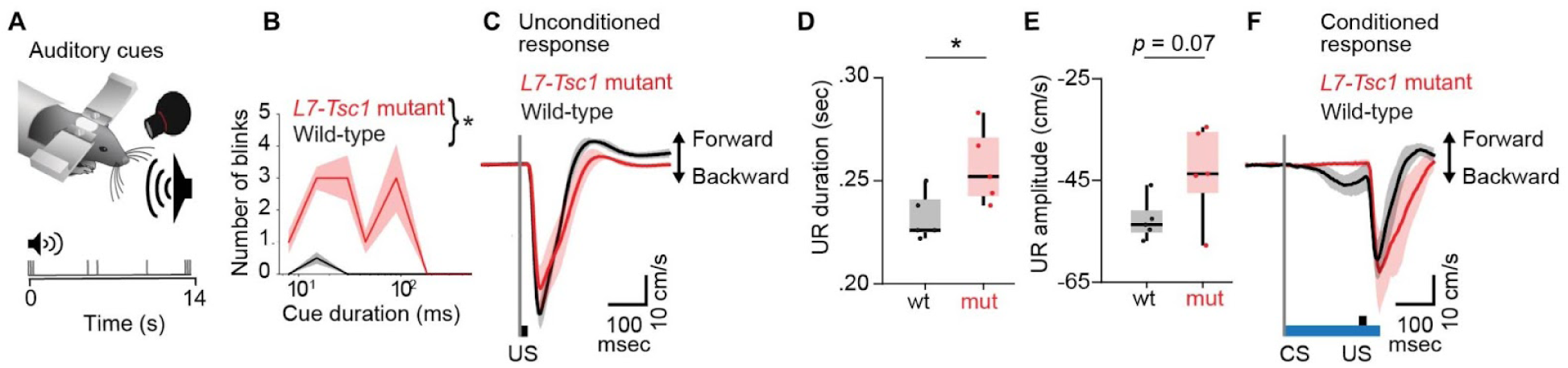
Increased sensory sensitivity in *L7-Tsc1* mice (A) Schematic of sensory sensitivity tests with auditory cues. (B) Median number of eye blinks in response to auditory cues of different durations for *L7-Tsc1* mutant mice (*n* = 16) and wild-type littermates (*n* = 7). A two-way ANOVA indicates an effect of genotype (F_(1)_ = 5.06, *p* = 0.026), but not of audio cue duration (F_(7)_ = 1.697, *p* = 0.11) or an interaction effect (F_(7)_ = 0.347, *p* = 0.93). (C) Unconditioned responses throughout sessions 1-5 in the DTSC task for *L7-Tsc1* mutant animals (red) and control animals (black). (D) *L7-Tsc1* mutant mice have an increased duration of the unconditioned response (UR) in the DTSC task compared to wild-type controls (F_(1)_ = 6.4, *p* = 0.04). (E) There is a trend toward larger amplitude of the unconditioned response (UR) in the DTSC task between *L7-Tsc1* mutant mice and wild-type controls (F_(1)_ = 4.5, *p* = 0.07). (F) Conditioned responses in session 5 in the DTSC task for *L7-Tsc1* mutant animals (red) and control animals (black). Shaded areas indicate the estimated s.e.m. using median absolute deviation.

**Figure S6.**
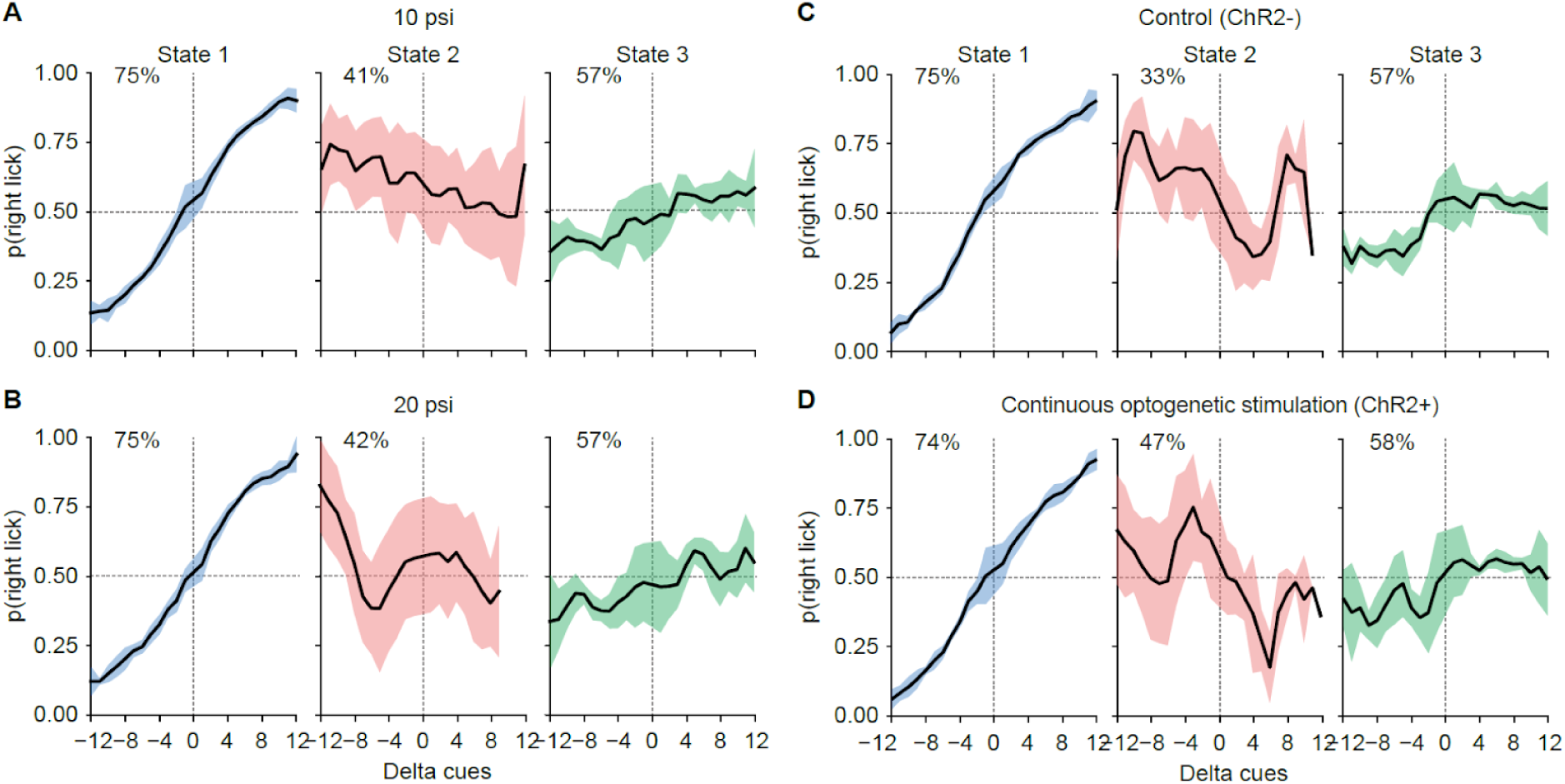
Performance and state occupancy for animals receiving stronger whisker puffs or optogenetic stimulation of Purkinje cells in crus I throughout the entire cue period and delay period (A-D) Psychometric curves for the three states, averaged across all mice receiving airpuffs to the whiskers of regular intensity (10 psi, A) or higher intensity (20 psi, B) or the normal task version for mice not expressing opsin (ChR2-, C) or for mice expressing opsin (ChR2+, D) and receiving continuous optogenetic stimulation to Purkinje cells in crus I during the cue period, delay period, and first lick. In the top left of each plot is the percentage correct over all trials in that state. Shaded areas represent one standard deviation.

**Figure S7.**
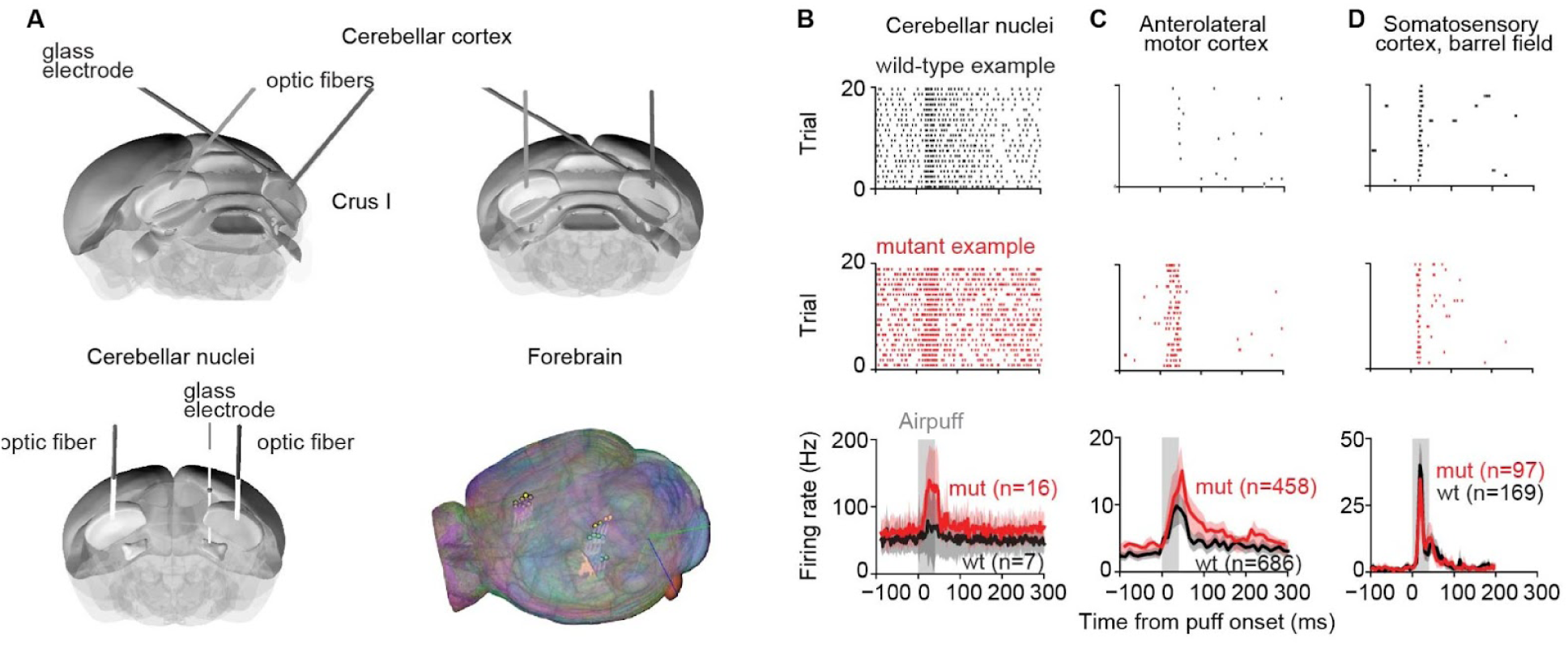
Increased responses to whisker puffs in *L7-Tsc1* mice in cerebellar nuclei and anterolateral motor cortex, but not somatosensory cortex (A) Orientation of the glass electrode during recordings in cerebellar cortex (top) and cerebellar nuclei (bottom left) combined with bilateral optogenetic stimulation of crus I. Bottom right: Examples of Neuronexus probe location in the forebrain for acute recordings in naive mice. (B) Example raster plots of cerebellar nuclei cells during 20 trials from *L7-Tsc1* mutants (middle) or their wild-type littermates (top), and average firing rates (bottom) in response to an airpuff to the whiskers (data from the same 4 *L7-Tsc1* mutants and 5 wild-type mice as in Figure 5I). There was a trend towards a significant difference for the AUC of the 100 ms post-puff minus the pre-puff firing rate (in spikes per 4 ms bin) for each unit (*L7-Tsc1* mutant mice: mean AUC = 12.0±9.8 (spikes/4ms bin × 100ms), wild-type littermates: mean AUC = 3.3±8.0 (spikes/4ms bin × 100ms), *t*_(22)_ = −1.97, *p* = 0.06, two-sided Student’s t-test). (C) Same as B, but for anterolateral motor cortex. There was a significant difference for the AUC of the 100 ms post-puff minus the pre-puff firing rate for each unit (*L7-Tsc1* mutant mice: mean AUC = 58.7±312.8, wild-type littermates: mean AUC = 26.7±221.1, *t*_(1143)_ = −2.02, *p* = 0.04, two-sided Student’s t-test). Baseline firing rates were not statistically different (*L7-Tsc1* mutant mice: *n* = 458 cells from 4 mice, mean = 1.26 Hz, wild-type littermates: *n* = 686 cells from 5 mice, mean = 1.14 Hz, *t*_(1143)_ = −1.04, *p* = 0.30, two-sided Student’s t-test). (D) Same as B and C, but for primary somatosensory cortex. There was no difference for the AUC of the 100 ms post-puff minus the pre-puff firing rate for each unit (*L7-Tsc1* mutant mice: mean AUC = 638±766, wild-type littermates: mean AUC = 694±1048, *t*_(265)_ = 0.45, *p* = 0.65, two-sided Student’s t-test). Shaded areas represent 95% confidence intervals.

**Figure S8.**
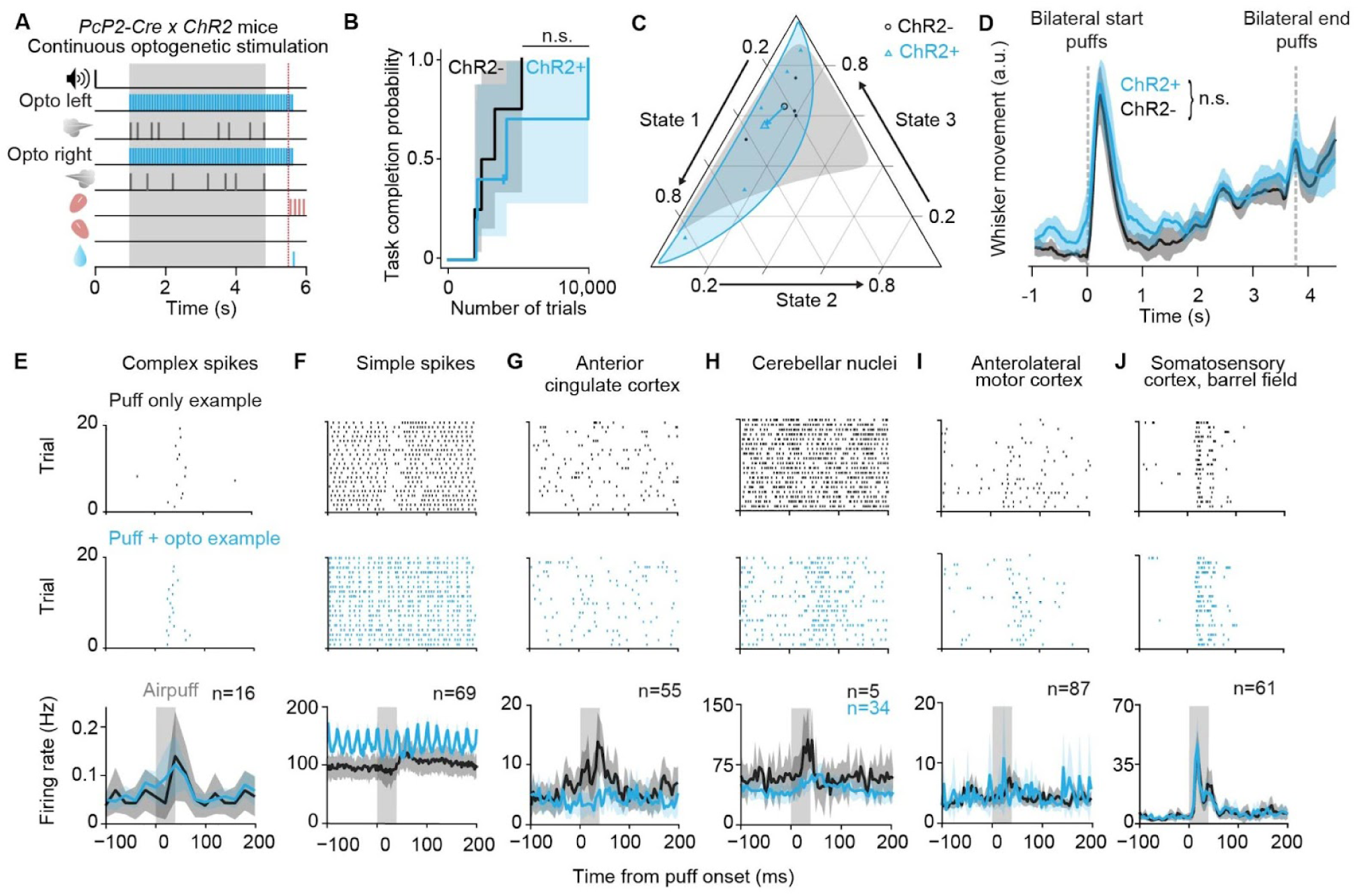
No effect of continuous optogenetic activation of Purkinje cells in crus I on learning of the evidence-accumulation task (A) Schematic of the evidence-accumulation task with continuous bilateral optogenetic activation of crus I. (B) Kaplan-Meier estimator of task completion probability for *PcP2-Cre* × *ChR2* mice with continuous bilateral optogenetic activation of crus I throughout the evidence-accumulation task (*n* = 5, median 4210 trials) compared to wild-type littermates (*n* = 4, median 2534 trials, *χ*^2^_(8)_ = 0.31, *p* = 0.33, log-rank test). (C) Ternary plot for the composition of latent states with 95% confidence bands (shaded regions) around the compositional mean (open symbols) for mice with continuous bilateral optogenetic activation of Crus I or controls. Analysis of variance on the log ratio state 2/state 1 yielded a significant effect for training level (F_(1)_ = 30.2, *p* < 0.001) but not for group (F_(1)_ = 2.9, *p* = 0.11). Analysis of variance on the log ratio state 3/state 1 yielded again a significant effect for training level (F_(1)_ = 6.4, *p* = 0.03) but not for group (F_(1)_ = 0.002, *p* = 0.96). The arrow indicates the shift from the average of the control group to the average of the group receiving continuous optogenetic stimulation. (D) Whisker movement during the evidence-accumulation task in response to bilateral whisker puffs at the start and end of each trial. Shaded areas include 95% confidence intervals. (E) Example raster plots of Purkinje cell complex spikes during 20 trials with only a whisker puff (top) or with a whisker puff paired with discounting optogenetic stimulation (middle), and average firing rates in response to an airpuff to the whiskers with or without paired optogenetic stimulation of Purkinje cells. (F) Same as E, but for Purkinje cell simple spikes. (G) Same as E, but for anterior cingulate cortex. (H) Same as E, but for cerebellar nuclei cells. (I) Same as E, but for anterolateral motor cortex. (J) Same as E, but for the barrel field of the primary somatosensory cortex. Shaded areas represent 95% confidence intervals.

**Figure S9.**
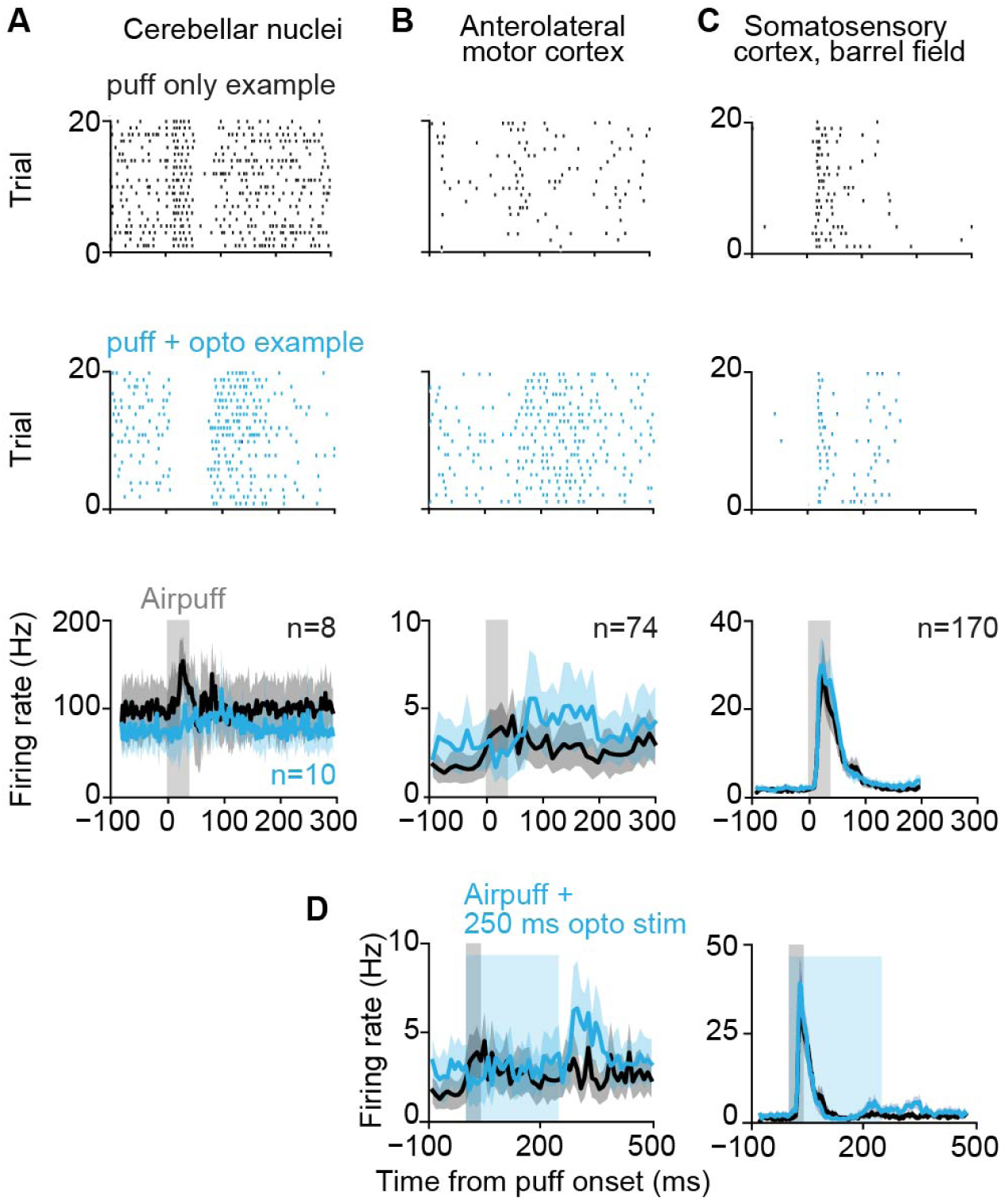
Altered responses to whisker puff and cue-locked optogenetic stimulation of Purkinje cells in crus I in cerebellar nuclei and anterolateral motor cortex, but not somatosensory cortex (A) Example raster plots of cerebellar nuclei cells during 20 trials with only a whisker puff (top) or with a whisker puff paired with optogenetic stimulation (middle), and average firing rates (bottom) in response to an airpuff to the whiskers with or without paired optogenetic stimulation of Purkinje cells in naive mice. Optogenetic stimulation significantly decreased the AUC 50 ms after stimulus offset (airpuff only: mean AUC = 17±7, airpuff combined with optogenetic stimulation: mean AUC = 10±6, *t*_(17)_ = 2.3, *p* = 0.03, two-sided Student’s t-test). (B) Same as A, but for anterolateral motor cortex. Optogenetic stimulation significantly increased the AUC 50 ms after stimulus offset (airpuff only: mean AUC = 38±66, airpuff combined with optogenetic stimulation: mean AUC = 61±102, *t*_(73)_ = −3.6, *p* = 0.004, paired t-test). (C) Same as A, but for the barrel field of the primary somatosensory cortex. Optogenetic stimulation did not have an effect on the AUC 50 ms after stimulus offset (airpuff only: mean AUC = 410±604, airpuff combined with optogenetic stimulation: mean AUC = 455±561, *t*_(169)_ = −1.3, *p* = 0.20, paired t-test). (D) Same as the bottom plots in B and C, but now with a longer duration (250 ms instead of 40 ms) of the optogenetic stimulation. Again, optogenetic stimulation significantly increased the AUC 50 ms after the end of the 250 ms long optogenetic stimulation (airpuff only: mean AUC = 38±66, airpuff combined with optogenetic stimulation: mean AUC = 62±118, *t*_(73)_ = −3.8, *p* = 0.0001, paired t-test), but not 50 ms after the end of the 40 ms long airpuff (airpuff only: mean AUC = 36±75, airpuff combined with optogenetic stimulation: mean AUC = 50±135, *t*_(73)_ = −1.7, *p* = 0.08, paired t-test). In S1, optogenetic stimulation also had an effect on the AUC 50 ms after the end of the 250 ms long optogenetic stimulation (airpuff only: mean AUC = 93±182, airpuff combined with optogenetic stimulation: mean AUC = 138±210, *t*_(169)_ = −2.8, *p* = 0.006, paired t-test), but not 50 ms after the end of the 40 ms long airpuff (airpuff only: mean AUC = 410±604, airpuff combined with optogenetic stimulation: mean AUC = 362±614, *t*_(169)_ = 1.3, *p* = 0.21, paired t-test). Shaded areas represent 95% confidence intervals.

**Figure S10.**
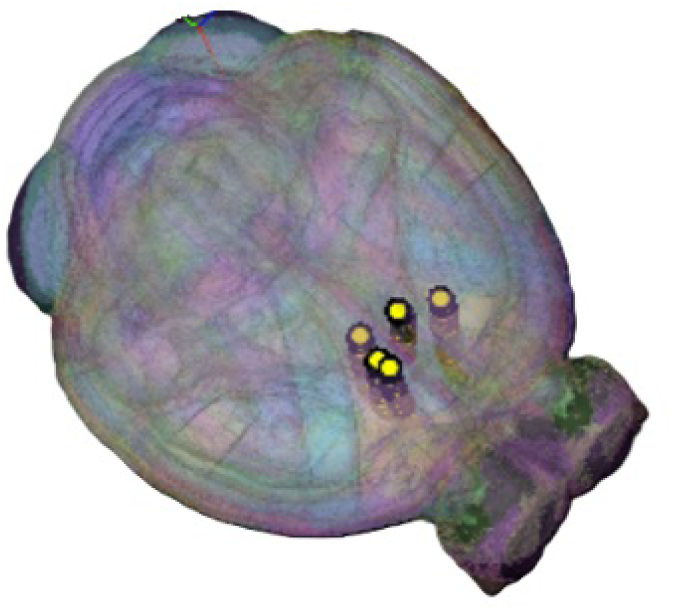
Recording location of Neuropixels probes in forebrain Visualization of the location of five Neuropixels probes targeted at anterior cingulate cortex for recording with simultaneous optogenetic stimulation of crus I while animals are performing the evidence accumulation task. Made with Neuroglancer (https://github.com/google/neuroglancer).

**Supplementary Table 1.** Mice progress through eight different levels during learning of the evidence-accumulation decision-making task.

| Level | 0 | 1 | 2 | 3 | 4 | 5 | 6 | 7 |
| --- | --- | --- | --- | --- | --- | --- | --- | --- |
| **Audio cue, 1s before cue period onset** | Yes | Yes | Yes | Yes | Yes | Yes | Yes | Yes |
| **Bilateral puffs at start** | Yes | Yes | Yes | Yes | Yes | Yes | Yes | Yes |
| **Cue period duration (s)** | 1 | 1 | 1 | 2.0, 2.8, or 3.8 | 3.8, or 1.5 | 3.8, or 1.5 | 3.8, or 1.5 | 3.8, or 1.5 |
| **Contralateral distractor puffs** | No | No | No | No | No | No | Yes, 1:9 | Yes, 1:4 |
| **Bilateral puffs at end** | No | No | Yes | Yes | Yes | Yes | Yes | Yes |
| **Delay** | 200 ms | 200 ms | 200 ms | 200 ms | 500 ms | 800 ms | 800 ms | 800 ms |
| **Guide puffs (2.5 Hz) until animal licks** | No | Yes | Yes | No | No | No | No | No |
| **Need to lick on correct side for reward** | No (both sides rewarded) | Yes | Yes | Yes | Yes | Yes | Yes | Yes |
| **Does first lick need to be correct** | No | No | Yes | Yes | Yes | Yes | Yes | Yes |
| **Error trials punished with tone and timeout** | No | No | Yes | Yes | Yes | Yes | Yes | Yes |
| **Requirements to proceed to next level** | 15 consecutive rewards | at least 100 trials & 55% correct in window of 40 trials | at least 200 trials & 80% correct in window of 50 trials | at least 100 trials & 75% correct in window of 40 trials | at least 100 trials & 80% correct in window of 40 trials | at least 25 trials & 80% correct in window of 24 trials | at least 250 trials & 75% correct in window of 40 trials | <75% correct is acceptable |

**Supplementary Table 2.**
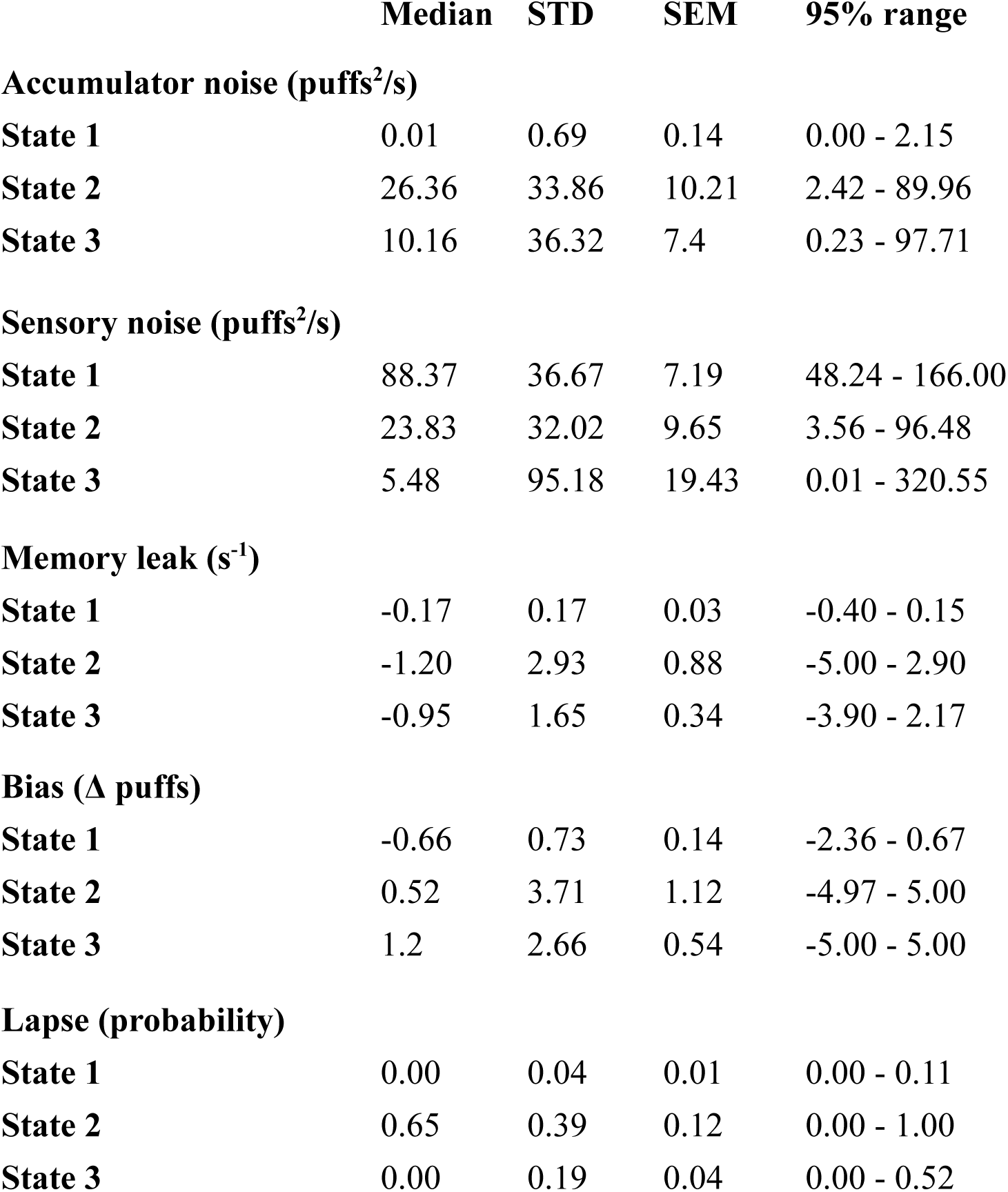
Best-fit drift diffusion model parameters for the three different latent behavioral states.

